# Across functional boundaries: making non-neutralizing antibodies to neutralize HIV-1 and mediate Fc-mediated effector killing of infected cells

**DOI:** 10.1101/2021.08.13.456231

**Authors:** Jonathan Richard, Dung N. Nguyen, William D. Tolbert, Romain Gasser, Shilei Ding, Dani Vézina, Shang Yu Gong, Jérémie Prévost, Gabrielle Gendron-Lepage, Halima Medjahed, Suneetha Gottumukkala, Andrés Finzi, Marzena Pazgier

**Author notes:** Jonathan Richard and Dung N. Nguyen equally contributed to this manuscript. The author order has been determined based on seniority. Corresponding authors To whom correspondence should be addressed, Centre de Recherche du CHUM, 900 St-Denis, Tour Viger, Montreal, QC, H2X 0A9, Canada, Uniformed Services University of the Health Sciences, 4301 Jones Bridge Road, Bethesda, MD 20814-4712.

## Abstract

In HIV-1 infection, many antibodies (Abs) are elicited to Envelope (Env) epitopes that are conformationally masked in the native trimer and are only available for antibody recognition after the trimer binds host cell CD4. Among these are epitopes within the Co-Receptor Binding Site (CoRBS) and the constant region 1 and 2 (C1-C2 or Cluster A region). In particular, C1-C2 epitopes map to the gp120 face interacting with gp41 in the native, ‘closed’ Env trimer present on HIV-1 virions or expressed on HIV-1 infected cells. Antibodies targeting this region are therefore non-neutralizing and their potential as mediators of antibody depended cellular cytoxicity (ADCC) of HIV-1 infected cells diminished by a lack of available binding targets. Here we present the design of Ab-CD4 chimeric proteins that consist of the Ab-IgG1 of a CoRBS or Cluster A specificity to the extracellular domain 1 and 2 of human CD4. Our Ab-CD4 hybrids induce potent ADCC against infected primary CD4+ T cells and neutralize tier 1 and 2 HIV-1 viruses. Furthermore, competition binding experiments reveal that the observed biological activities rely on both the antibody and CD4 moieties confirming their cooperativity in triggering conformational rearrangements of Env. Our data indicate the utility of these Ab-CD4 hybrids as antibody therapeutics effective in eliminating HIV-1 through the combined mechanisms of neutralization and ADCC. This is also the first report of single-chain-Ab-based molecules capable of opening ‘closed’ Env trimers on HIV-1 particles/infected cells to expose the Cluster A region and activate ADCC and neutralization against these non-neutralizing targets.

**Importance:** Highly conserved epitopes within the co-receptor binding site (CoRBS) and constant region 1 and 2 (C1C2 or Cluster A) are only available for antibody recognition after the HIV-1 Env trimer binds host cell CD4, therefore they are not accessible on virions and infected cells where the expression of CD4 is downregulated. Here we have developed new antibody fusion molecules in which domains 1 and 2 of soluble human CD4 are linked with monoclonal antibodies of either the CoRBS or Cluster A specificity. We optimized the conjugation sites and linker lengths to allow each of these novel bispecific fusion molecules to recognize native “closed” Env trimers and induce the structural rearrangements required for exposure of the epitopes for antibody binding. Our *in vitro* functional testing shows that our Ab-CD4 molecules can efficiently target and eliminate HIV-1 infected cells through antibody depended cellular cytoxicity (ADCC) and inactivate HIV-1 virus through neutralization.

## Introduction

The envelope glycoprotein trimer (Env) plays a crucial role in HIV-1 virus attachment and entry into host cells. The mature Env trimer is comprised of non-covalently associated gp120-gp41 heterodimers that are formed by furin cleavage of a gp160 precursor(1). To initiate the viral entry process, the outer gp120 protomer of the trimer binds the receptor CD4 on the host cell surface. Post CD4-binding, Env undergoes conformational changes that leads to the formation of the co-receptor binding site (CoRBS) and engagement of CCR5 or CXCR4, the two known HIV-1 co-receptors (2–9). Post-co-receptor binding, additional structural rearrangements within Env occurs that leads to the formation of a six-helix bundle from the helical heptad repeat HR1 and HR2 segments of the gp41 ectodomain to drive fusion of viral and target cell membranes(10, 11). The trimeric HIV-1 Env is also the sole viral protein present on the surface of virions and HIV-1-infected cells and thus, it represents the major antibody-targeted HIV-1 antigen. Env presentation to the host immune system elicits antibody (Ab) responses against many diverse Env sites. These Abs can impact HIV-1 through various mechanisms including direct virus neutralization or Fc-effector activities such as antibody-dependent cellular cytotoxicity (ADCC) of infected cells. In HIV-1 infection, a number of elicited antibodies target epitopes which are occluded in the unliganded Env trimer present at the surface of infected cells or infectious viral particles. These Abs usually lack direct neutralization activity and therefore are referred to as non-neutralizing (nnAbs)(12, 13). Importantly, some Env sites recognized by nnAbs map to highly conserved regions of Env so their potential utility as targets for protective humoral responses/antibody therapeutics is deemed high, assuming one can overcome the obstacle associated with their lack of exposure.

Among the most prominent targets for nnAbs are epitopes that become available post-CD4 binding (CD4-induced or CD4i) when Env transitions from its unliganded “closed” conformation (State 1) to an “open” CD4-bound conformation (State 3)(14). The two epitopes that become available sequentially in this process are 1) CoRBS and 2) Cluster A, epitopes within the mobile layer 1 and 2 of the highly conserved constant region 1 and 2 (C1C2) of the gp120 inner domain(12, 13, 15–23). Whereas the CoRBS is localized at the surface of the Env trimer (mapping to the outer domain of gp120, proximal to the CD4 binding site), the Cluster A epitopes map to the interior of HIV Env trimer at the gp41-gp120 interface and are directly involved in inter-promoter contacts that stabilize the trimer. The exposure of Cluster A region requires significant structural rearrangements of gp120 and gp41 subunits, something that occurs late in the entry process as a consequence of CD4-induced changes in Env(12). Therefore, all known Cluster A Abs uniformly lack direct neutralizing activities(19). Interestingly, CD4i nnAbs are frequently elicited in HIV-1-infected individuals and are capable of mediating potent ADCC against CD4i targets(16, 17, 24, 25). Unfortunately, their potential as ADCC mediators is greatly diminished by the fact that cells infected with primary HIV-1 isolates express Env in a ‘closed’ conformation in which CD4i targets are not accessible for Ab recognition(14, 26, 27). We and others have shown that HIV-1 has evolved to preserve Env in a “closed” conformation and limit CD4 interaction to prevent exposure of these ADCC targets that are otherwise recognized by easily-elicited Abs present in the sera of HIV-1-infected individuals(16, 17, 24, 27–29). Internalization of Env(30, 31), CD4 downregulation via the HIV-1 accessory proteins Nef and Vpu (16, 17), and Vpu-mediated antagonism of the host restriction factor BST-2(16, 17, 32, 33) were found to be contributing resistance mechanisms protecting infected cells or viral particles(34) from CD4i nnAbs. This could explain why the majority of circulating HIV strains express functional Nef and Vpu proteins.

To overcome these protective mechanisms, soluble CD4 (sCD4) or small CD4-mimetic (CD4mc) compounds have been used to modulate Env conformation and expose CD4i epitopes, mostly within the CoRBS for direct neutralizing activity(35). Treatment with CD4mc was shown to sensitize primary strains of HIV-1 to neutralization by CD4i Abs(36) and sCD4-17b bifunctional proteins(37, 38) or eCD4-Ig variants incorporating CD4 IgG and a CoRBS-specific sulfo-peptide into a single construct(39) were developed to create molecules with exceptional breadth and neutralization potency against HIV-1 strains. Interestingly, sCD4 and CD4mc have also been used to expose CD4i Env epitopes on infected cells despite Nef and Vpu expression, and effectively sensitize them to ADCC by CD4i nnAbs present in sera, breastmilk and mucosa of HIV-1-infected individuals(40). However, the mechanism of sensitization of infected cells to ADCC requires a sequential opening of the Env trimer and depends on the cooperation of sCD4/CD4mc in addition to CoRBS and Cluster A nnAbs(21, 23, 29). In this scenario, a mixture of sCD4 or CD4mc, a CoRBS Ab and a Cluster A Ab is required to trigger the asymmetric ADCC vulnerable Env conformation referred to as State 2A, an intermediate between State1 and 3 in the opening of the Env trimer(21, 23, 41).

Here, we thought to develop single antibody-based molecules that could combine these elements to eliminate HIV-1 infected cells by ADCC. We engineered constructs in which a CoRBS or Cluster A nnAb IgG is linked by a (G_4_S)_6_(G_4_T)_2_-linker to sCD4 (d1d2 domain, residues 26-208) to generate a single chain molecule referred to as an Ab-CD4. Functional analyses fully confirmed this approach as Ab-CD4 proteins can efficiently target and eliminate HIV-1-infected cells by ADCC through a mechanism involving the binding of both the nnAb and CD4 moieties. Furthermore, both the CoRBS and the Cluster A Ab-CD4 molecules showed direct neutralization activity against tier 1 and 2 HIV-1 viruses. Altogether, these new Ab-CD4 molecules are capable of impacting HIV-1 through a coordinated mechanism of direct neutralization of virions and ADCC against infected cells. This is to our knowledge also the first successful report of activation of direct neutralization for Abs of Cluster A region traditionally known for having no neutralizing activities.

## Results

### Design and expression of single chain Ab-CD4

The single chain Ab-CD4 molecules, consist of a CoRBS or Cluster A specific nnAb linked to the C-terminus of sCD4 (domain 1 and 2, residues 26-208 of extracellular domain of human CD4) via a flexible 40 amino acid (aa) –(-(Gly_4_--Ser)_6_-(Gly_4_--Thr)_2_) linker (Fig. 1). The linker is attached to the N-terminus of the heavy chain (IgGH) of the mAb IgG1 resulting in molecule in which 2 sCD4 domains are attached to single IgG1. For each epitope specificity, 2 antibodies were tested as the IgG1 arm of the Ab-CD4 molecule. A32, the prototype antibody of the Cluster A region (19), and N5-i5, an Ab isolated from an HIV-1 infected individual capable of potent ADCC against CD4i Env targets(18, 19), were selected to represent the Cluster A region. A32 and N5-i5 recognize largely overlapping epitopes that map to the highly conserved C1-C2 portion of the Cluster A region of CD4-triggered gp120 (18, 42) (Fig.1a). For the CoRBS-specific antibodies, we selected 17b, an Ab recognizing a conserved epitope within the bridging sheet of the CoRBS (43, 44) and X5, an Ab combining the elements of the highly conserved bridging sheet of the CoRBS with elements of the V3 loop stem(45, 46). The length of the (G_4_S)_6_-(G_4_T)_2_-linker was selected based upon structural information of the antibody epitope within the gp120 promoter in relation to the position of the CD4 binding site in the Env trimer (Fig. 1a). The selected linker length is compatible with the binding of the Fab arm and sCD4 to the same gp120 promoter in the trimer as well as to possibly crosslink two adjacent gp120 protomers within the same trimer. Our selected linker length is similar to what was used to develop conjugates of a single chain variable fragment (scFv) of 17b and sCD4 that showed HIV-1 neutralizing activity(37). ScFv conjugates with long linkers (35-40 aa residues) displayed stronger neutralizing activity as compared to the same constructs with a shorter linker (5-20 aa residues). Therefore, we designed our Ab-CD4s with a flexible 40-aa linker (six repeats of the G_4_S motif and two repeats of the G_4_T motif).

**Fig. 1.**
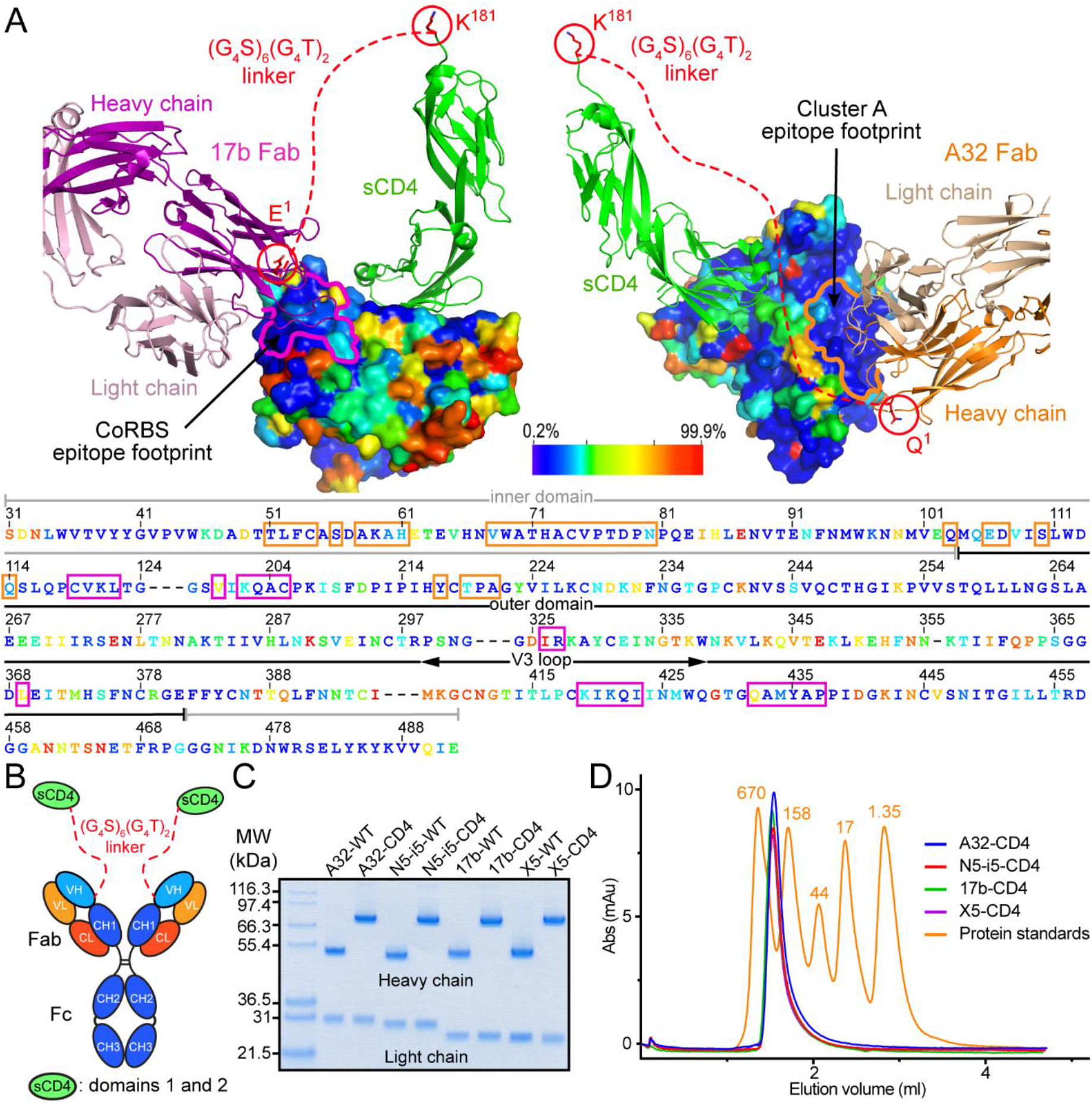
Generation of the Ab-CD4 hybrid proteins. (**A**) Cartoon representation showing the N-terminus residues E^1^ and Q^1^ of the heavy chain of 17b mAb and A32 mAb, respectively, as conjugation positions to the C-terminus of human receptor CD4 (domains 1 and 2). Conjugation positions are circled in red. The flexible (G_4_S)_6_(G_4_T)_2_ peptide linker is represented as the red dashed line. The molecular surface is displayed over gp120 and colored based on the sequence conservation. Residues differed from the Hxbc2 sequence between 0.2-7% are colored with dark blue and residues differed between 87-99.9% are colored in red. 17b Fab and A32 Fab are shown as ribbon diagram. The heavy chain and light chain of mAb 17b are colored in magenta and light pink, respectively. The heavy chain and light chain of mAb A32 are colored in orange and light orange, respectively. The CoRBS and cluster A epitope footprints are highlighted in purple and orange on the gp120 surface, respectively. The sequence of gp120, including variable region, is shown based on the conservation of each residue. The CoRBS and cluster A epitope footprints are highlighted with purple line and orange line, respectively. (**B**) Schematic drawing of Ab-CD4 molecules. Domains 1 and 2 of CD4 protein (green) were linked to the N-terminus of the antibody heavy chain through a 40 amino acid linker (6 repeats of G_4_S and 2 repeats of G_4_T motifs). VH = variable heavy (light blue), VL = variable light (light orange), CH = constant heavy (blue) and CL = constant light (orange). (**C**) SDS-PAGE gel of four wild-type antibodies (WT-Abs) and four Ab-CD4 molecules under reducing condition after size exclusion chromatography. The heavy chains of WT-Abs migrate at ~50 kDa and the heavy chains of Ab-CD4 migrate at ~75 kDa. A NuPAGE™ 4-12% Bis-Tris protein gel (Invitrogen) was used to separate the protein bands with Mark 12™ Unstained Protein Standard (Invitrogen) in the first lane for reference. (**D**) Size exclusion chromatography (SEC) of all four purified Ab-CD4 molecules using Superdex 200 Increase 5/150 GL column as compared to the gel filtration protein standards (BioRad, #1511901).

Ab-CD4 molecules were produced in mammalian Expi293F cells. First, we developed an Expi293F cell line stably expressing domains 1 and 2 of CD4 fused to the N-terminus of the heavy chain of the selected nnAb. We then transfected the plasmid containing the appropriate kappa or lambda light chain. The production of Ab-CD4s using this method typically yielded 6-19 mg of properly folded product per liter of culture. The typical purification protocol included Protein A affinity chromatography, followed by size exclusion chromatography (SEC). The calculated molecular weight of the Ab-CD4 bifunctional molecule is roughly 200 kDa as compared to 150 kDa for wild-type mAb. The SEC polishing step usually removed any protein aggregates or dimers. Fig. 1c and d shows the SDS-PAGE and SEC analyses, respectively, of the final Ab-CD4 preparations. Under the reducing condition of SDS-PAGE, all the mAbs, alone or linked to sCD4, were resolved at the expected molecular weights (Fig. 1c). The heavy chains of all Ab-CD4s migrate to form a band with a molecular weight of 75 kDa that corresponds to the sum of the heavy chain (~50kDa) plus the d1d2 domain of CD4 (~25kDa). In SEC analyses, one distinct peak corresponding to the Abs-CD4 was observed eluting earlier than a 158 kDa peak of mAb alone (Fig. 1d).

### Ab-CD4 single chain molecules recognize the unliganded Env present at the surface of HIV-1-infected cells

Ab-CD4 molecules were developed to recognize the ‘closed’ Env trimer available at the surface of HIV-1 infected cells to trigger and expose CD4i epitopes. Therefore, we first evaluated their binding capacity to unliganded cell-surface Env using a previously described cell-based ELISA (47, 48). Briefly, HOS cells were transfected with a plasmid encoding the HIV-1_JRFL_ Env. Two days later, transfected cells were incubated with Abs alone (17b, X5 and A32, N5-i5), Abs plus soluble CD4 (sCD4) mix or Ab-CD4s and binding was detected using HRP-conjugated secondary Abs. With agreement to the data available on exposure of CoRBS and Cluster A epitopes neither of nnAbs tested alone was able to recognize Env-expressing cells (Fig. 2a). When mixed with sCD4, the CoRBS specific nnAbs 17b and X5 but not Cluster A specific nnAbs A32 and N5-i5 recognized cell-surface Env. This is consistent with published information on Env conformations post sCD4 binding, where sCD4 is able to trigger the conformation changes required to expose the CoRBS but not the Cluster A region(21, 23, 29). However, Ab-CD4 single chain molecules consisting of the Ab arm of either of the two CD4i nnAb classes efficiently recognized Env-expressing cells. Interestingly, the binding levels of Cluster A Ab-CD4s were comparable to CoRBS Ab-CD4s specificities indicating the same level of epitope exposure. This is of particular interest as the Cluster A region maps to the Env trimer interior and requires significant structural rearrangements of the trimer post CD4-binding for exposure(19, 49). This also indicates that the requirement of cooperativity between CoRBS and Cluster A nnAbs for efficient recognition of unliganded cell-surface Env pretreated with sCD4/CD4mc(21, 50) can be overcome with a single chain Cluster A Ab-CD4. We further tested the capacity of the developed Ab-CD4s to recognize HIV-1-infected cells by flow cytometry (Fig. 2b-d). Activated primary CD4+ T cells were infected with the primary HIV-1 isolate JRFL and two days later, the infected cells were incubated with Ab-CD4 or nnAb. Antibody binding was measured using an Alexa fluor-647-conjugated secondary Abs. Consistent with the protective activities of Nef and Vpu proteins which limit cell-surface Env-CD4 interaction, infected primary CD4+ T cells were mainly resistant to recognition by mAbs targeting the CoRBS (17b and X5) or the Cluster A (N5-i5 and A32) regions when used alone (Fig. 2b,c). While addition of sCD4 significantly enhanced the capacity of the CoRBS Abs 17b and X5 to recognize HIV-1-infected cells, both CoRBS Ab-CD4 molecules (17b-CD4 and X5-CD4) directly recognized infected cells. Interestingly, the capacity of CoRBS Ab-CD4s to recognize HIV-1-infected cells was similar or superior to their unconjugated Abs counterpart. In contrast, Cluster A Abs failed to recognize infected cells either when used alone or in the presence of sCD4 and only Cluster A Ab-CD4s (N5-i5-CD4 and A32-CD4) efficiently bound to JRFL-infected cells. Importantly, all developed Ab-CD4 specifically recognized HIV-1-infected cells over autologous mock-infected cells (Fig. 2dd) and triggered Env conformational changes as indicated by increased binding of the anti-gp41 mAbs F240 upon Ab-CD4 interaction (Fig. S1). Since F240 targets a gp41 epitope that is only exposed late in the entry process post CD4(51) binding similar to CD4i Env epitopes, these data suggest that Ab-CD4 molecules ‘push’ the Env trimer into similar downstream conformational states. This conformation is likely ‘more open’ than that present when triggered by sCD4 alone or with sCD4 nnAb mix, as indicated by the more efficient F240 epitope exposure for Env treated with Ab-CD4 as compared to nnAb, sCD4 or sCD4 + nnAb mix.

**Fig. 2.**
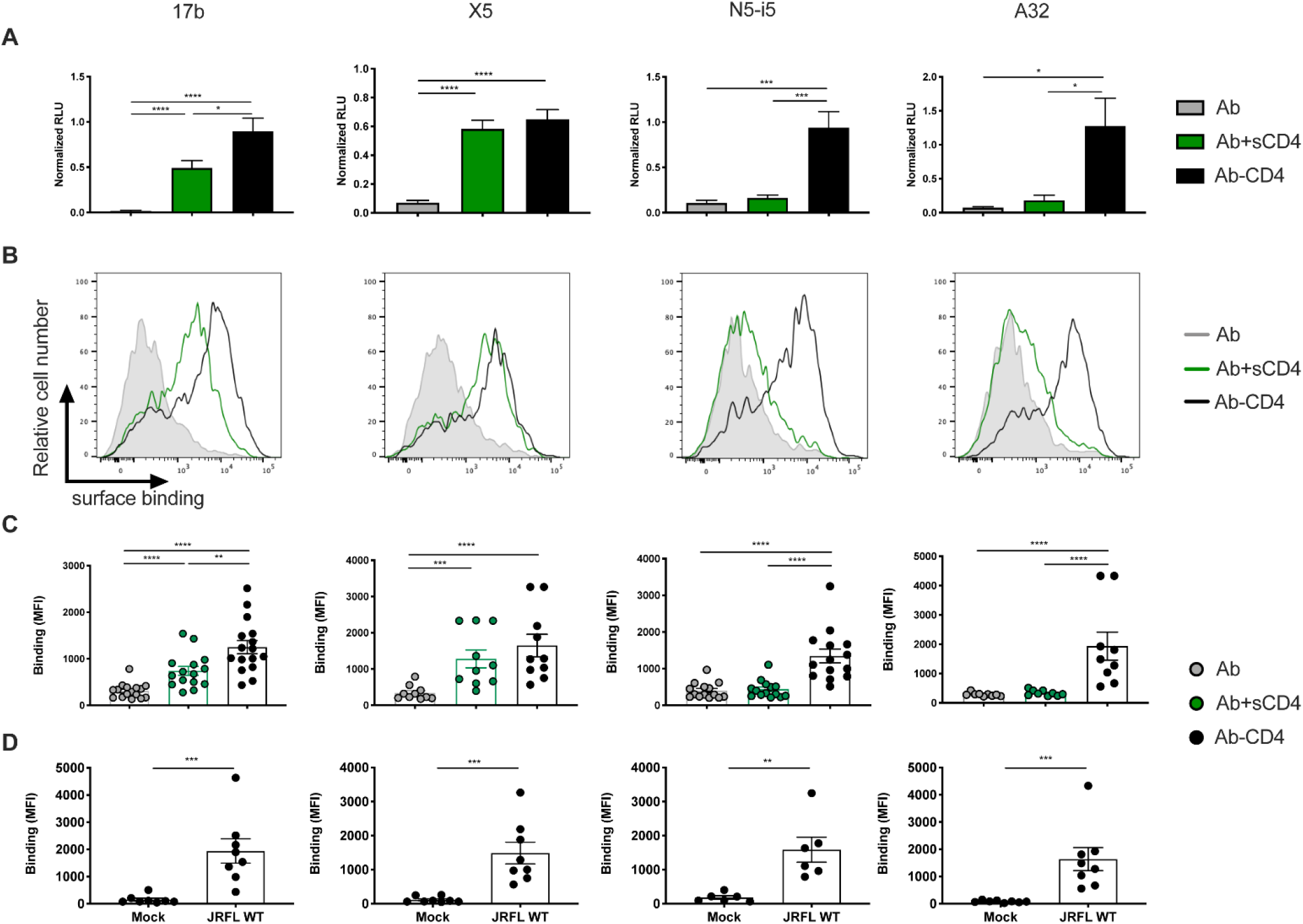
CoRBS Ab-CD4 and Cluster A Ab-CD4 directly recognize HIV-1-infected cells. (**A**) Recognition of cellular-expressed trimeric Env by indicated Ab-CD4 or Ab in the presence or absence of sCD4 (3 μg/ml) was evaluated by cell-based ELISA. Data shown represent mean RLU values ± SEM from at least 4 independent experiments performed in quadruplicate, with the signal obtained from cells transfected with an empty pcDNA3.1 plasmid (no Env) subtracted, normalized to Env levels as determined by bNAb 2G12. (b-d) Recognition of primary CD4+ T cells infected with HIV-1 JRFL or mock-infected by indicated Ab-CD4 or Ab (10 μg/ml) in the presence or absence of sCD4 (10 μg/ml) was evaluated by flow cytometry. (**B**) Histograms depicting representative staining. (**C**-**D**) The graphs shown represent the compiled median fluorescence intensities on the infected (p24+) cell population (infected cells) or mock-cell population. Error bars indicate means±SEM for at least 6 independent experiments. Statistical significance was tested using an unpaired t-test or a Mann-Whitney U test based on statistical normality (*, *P* < 0.05; **, *P* < 0.01, ***, *P* < 0.001; ****, *P* < 0.0001)

### Ab-CD4s eliminate HIV-1-infected cells through an ADCC mechanism

The efficient recognition of HIV-1 infected cells by Ab-CD4s prompted us to test if this translates into ADCC against infected cells. The ADCC activity was evaluated using a previously described flow cytometry-based ADCC assay(40) using primary CD4+ T cells infected with JRFL as target cells and autologous PBMCs as effector cells. The ADCC-mediated elimination of infected cells was determined by the loss of p24+ target cells upon treatment with Ab-CD4 or Ab and effector cells. Since Ab-CD4 showed similar biological activity within each class of nnAb, one fusion Ab from each class was tested (17b-CD4 and N5-i5-CD4). Previous studies showed that the CoRBS Ab 17b does not mediate efficient ADCC, even against cells with exposed CD4-bound Env, such as infected cells treated with CD4mc or cells infected with Nef and Vpu defective virus (21, 23, 24, 29). Accordingly, JRFL-infected cells were found to be resistant to ADCC mediated by 17b, even in the presence of sCD4 (Fig. 3). In contrast, an Ab-CD4 molecule 17b mediates ADCC against JRFL-infected cells (Fig. 3). This confirms that effective recognition of infected cells translated to ADCC. Most importantly we also observed a similar ADCC killing with the N5-i5-CD4, indicating effective triggering of effector cells and ADCC by single chain CD4 molecules targeting the Cluster A region. Although it is known that Cluster A Abs are capable of ADCC of cells exposing Env in the CD4-bound conformation(19–21, 24, 26, 29, 52), the infected cell recognition and ADCC is poor due to epitope occlusion(21, 23, 29). Consequently, primary CD4+ T cells infected with JRFL were found to be resistant to ADCC by N5-i5 alone or in combination with sCD4 (Fig. 3). These results suggest that both classes of Ab-CD4 can directly recognize and mediate ADCC of HIV-1-infected cells.

**Fig. 3.**
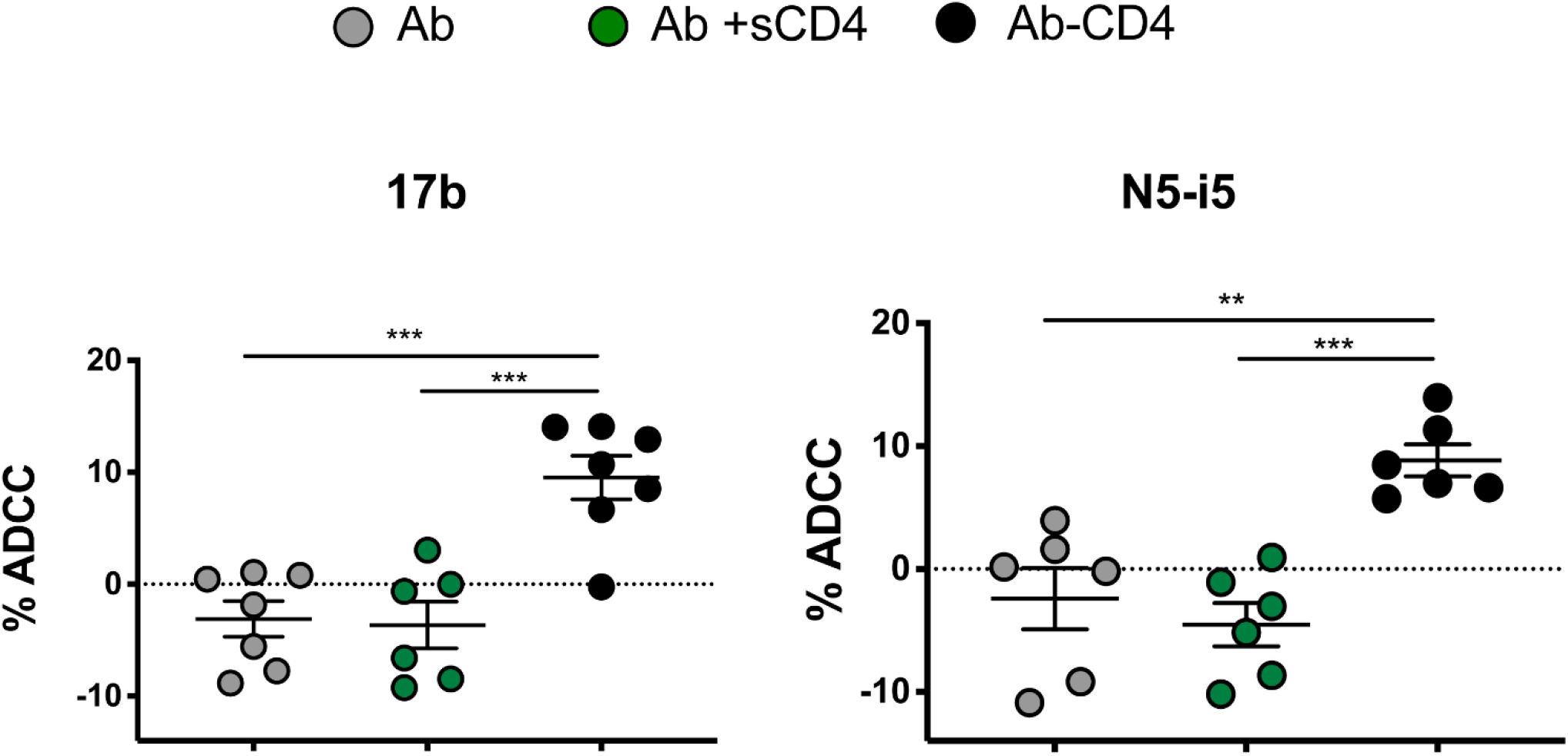
CoRBS Ab-CD4 and Cluster A Ab-CD4 mediate ADCC responses against HIV-1-infected cells. Primary CD4+ T cells infected with HIV-1 JRFL were used as targets and autologous PBMCs as effector cells in a FACS-based ADCC assay. The graphs shown represent the percentage of obtained in the presence of indicated Ab-CD4 or Ab in the presence or absence of sCD4. These results were obtained in at least 6 independent experiments. Statistical significance was tested using an unpaired t-test (**, *P* < 0.01, ***, *P* < 0.001)

### Ab-CD4s show neutralizing activity against tier 1 and 2 HIV-1 Env

While some Abs specific for CoRBS are capable of weak neutralization of Tier 1 viruses(53–55), the Cluster A Abs represent a group of canonical non-neutralizing Abs incapable of impacting virus through neutralization. Since Ab-CD4 molecules with an Ab arm of either CoRBS or Cluster A specificity bound to Env present on infected cells, we decided to verify if they could recognize the trimeric Env present on HIV-1 virions. We therefore used a previously described virus-capture assay(34) which measures binding of HIV-1 virions by mAbs immobilized on enzyme-linked immunosorbent assay (ELISA) plates. Viral particles were produced by co-transfecting HEK293T cells with the pNL4.3 Nef Luc Env-construct, a plasmid encoding the tier-2 HIV-1_JRFL_ Env and a plasmid encoding the G glycoprotein from the vesicular stomatitis virus (VSV-G). This generates a virus capable of a single round of infection. Virus containing supernatants were added to plates coated with Ab or Ab-CD4 and unbound virions removed by washing. Antibody-mediated retention of HIV-1 virions was assessed by addition of HEK293T cells. Infection of this CD4-negative cell-line is mediated by VSV-G and can be measured by luciferase activity 2 days post-infection. Env from primary HIV-1 isolates, such as JRFL, naturally adopt a closed conformation in which the CoRBS and the Cluster A regions are not exposed(14, 21). Accordingly, CoRBS Abs (17b and X5) and Cluster A Abs (N5-i5 and A32) failed to capture HIV-1 virions (Fig. 4a). Furthermore, in agreement with the pattern observed before for HIV-1 infected cells, we observed an enhanced capacity for CoRBS Abs to capture HIV-1 viral particles in presence of sCD4, but this was not the case for Cluster A Abs which were unable to retain virions either in the presence or absence of sCD4. For Cluster A Abs we observed virion capture only when the Ab was directly linked to CD4 confirming again that Cluster A-CD4 molecules are able to trigger conformational rearrangements of the Env trimer required for C1C2 epitope exposure (Fig. 4a). Notably in all cases, the retention of virions by Ab-CD4 tend to be superior to their unconjugated counterpart Ab +/− sCD4.

**Fig. 4.**
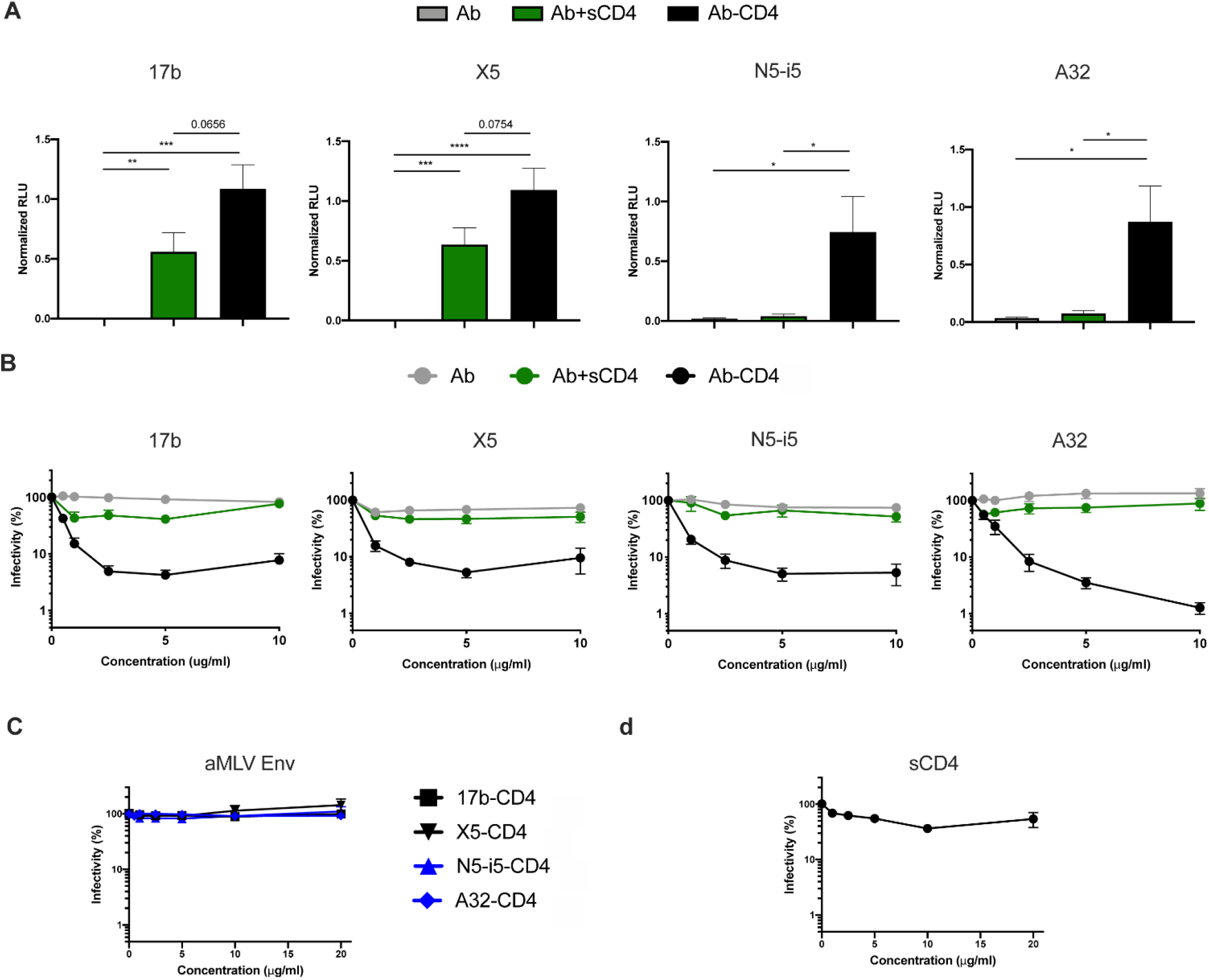
CD4-attachment enhances the capacity of CoRBS and Cluster A Abs to capture and neutralize HIV-1 viral particle. (**A**) The capacity of indicated Ab-CD4 or Ab to capture VSV-G pseudotyped viral particle expressing HIV-1_JRFL_ Env was assessed by virus-capture assay. Viral particles were added to plate coated with different Ab-CD4 or counterpart Ab in the presence or absence of sCD4. Free virions were washed away, and HEK293T cells were added to the wells. After 48 h, cells were lysed and luciferase activity was measured. Luciferase signals were normalized to those obtained with the 2G12 antibody. These results were obtained in 3 independent experiments. Error bars indicate mean ± standard error of the means (SEM). Statistical significance was calculated using unpaired t test or Mann-Whitney test based on normality test. (**B**-**D**) Viral particles pseudotyped with HIV-1_JRFL_ Env (B,D) or the A-MLV Env (C) were incubated with serial dilution of indicated Ab-CD4, Ab (+/− 5ug/ml of sCD4) or sCD4 before infecting TZM-bl cells. Luciferase activity in cell lysates was measured to determine the infectivity. Infectivity was normalized to 100% in the absence of the ligand. Data shown are representative of results from at least three experiments.

Finally, we evaluated whether the increased capacity of Ab-CD4s to capture viral particles translated into direct neutralizing activity (Fig. 4b). While CoRBS (17b or X5) and Cluster A (N5-i5 or A32) Abs alone failed to neutralize pseudoviruses bearing the Env from JRFL, both Ab-CD4 classes did (Fig. 4bb). This activity was specific to HIV-1 Env, since no neutralization was observed against pseudoviruses bearing A-MLV Env (Fig. 4cc). Importantly, the neutralizing activity of Ab-CD4 was superior to sCD4 alone or in combination with nnAbs, confirming that these hybrid molecules represent an improvement over the mixture of both moieties together (Fig 4b,d). We then tested the neutralizing capacity of these molecules against a panel of virus or pseudoviruses bearing HIV-1 Env from tier 1 and 2 strains. Consistent with the conserved nature of the epitopes targeted by the two classes of Ab-CD4 (Fig. 1a), they showed neutralization activities against viruses or pseudoviruses bearing HIV-1 Env from clades A, B, C and CRF01_AE strains (Fig. 5).

**Fig. 5.**
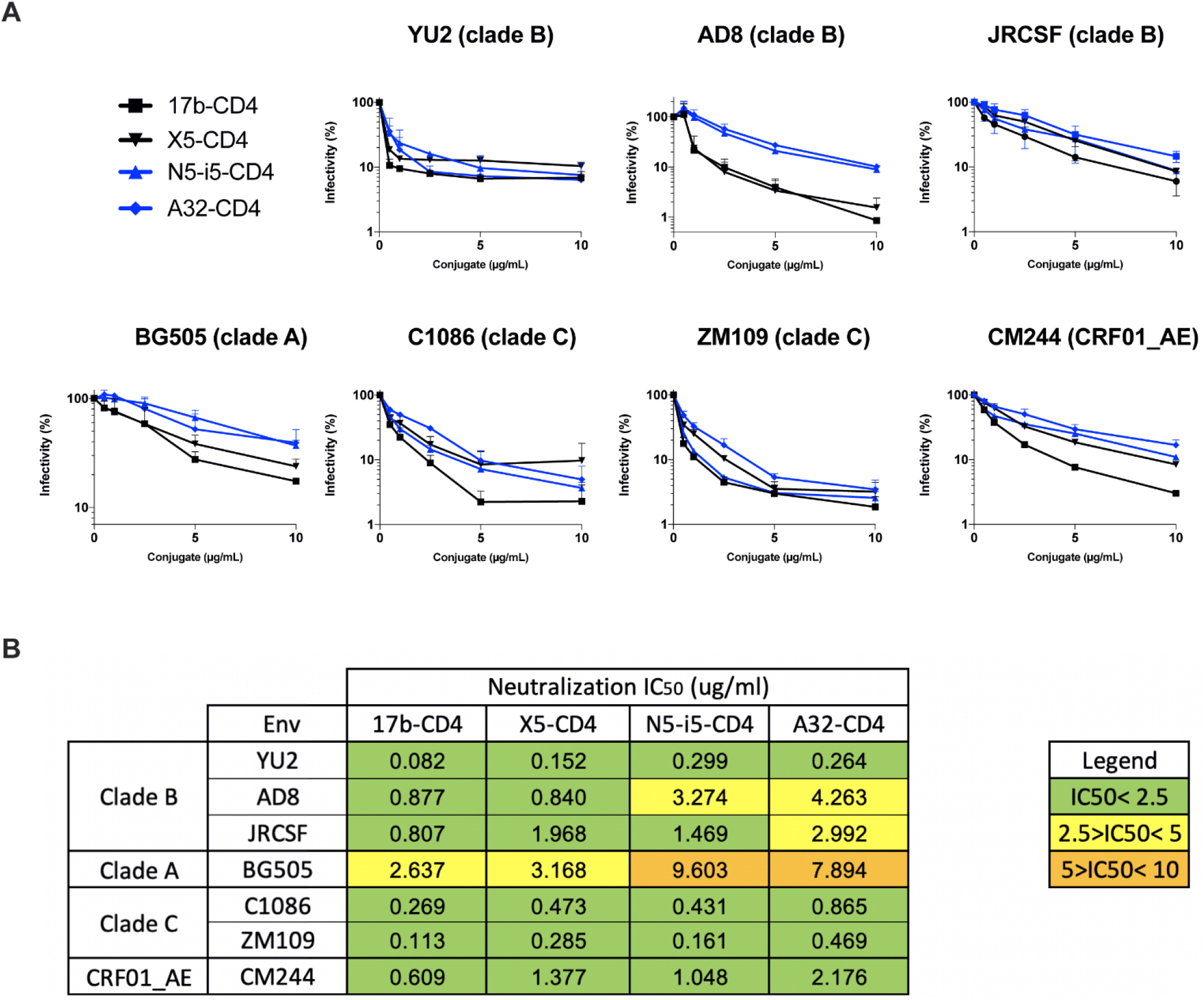
CoRBS Ab-CD4 and Cluster A Ab-CD4 neutralize viruses from different clades. (**A**) Viral particles from HIV-1 AD8, JR-CSF or pseudotyped with Envs HIV-1_YU2_, HIV-1_BG505_, HIV-1_C1086_, HIV-1_ZM109_ or HIV-1_CM244_ were incubated with serial dilution of indicated Ab-CD4 before infecting TZM-bl cells. Luciferase activity in cell lysates was measured to determine the infectivity. Infectivity was normalized to 100% in the absence of Ab-CD4s. Data shown are representative of two independent experiments. The error bars represent means±SEM. Neutralization half-maximal inhibitory concentration of the different Ab-CD4 for each virus are summarized in (**B**).

### The biological activities of Ab-CD4s rely on both Ab and CD4 moieties interaction with Env

Next, we aimed to investigate the mechanism of interaction of our Ab-CD4s with the Env trimer, in particular, to determine if both the CD4 and the Ab arms are engaged in binding to Env on HIV-1-infected cells/viral particles and are involved in the ADCC/neutralization. To assess the specific role of the CD4 moiety, we introduced a D368R mutation into the CD4 binding site of HIV-1_JRFL_ Env. This mutation was shown to abrogate the Env-CD4 interaction (16, 41, 50, 56)(Fig. S2). As presented in fig. 6a, introduction of this mutation totally inhibited the capacity of both classes of Ab-CD4 to recognize cell-surface expressed HIV-1_JRFL_ Env. This confirms that the sCD4 moiety of these Ab-CD4 is required to target the unbound “closed” Env. To assess the specific role of the antibody moiety in the biological activities of our developed Ab-CD4, competition experiments were done using two different gp120 probes. We first used a CD4-bound stabilized gp120 core protein, lacking variable regions V1, V2, V3, and V5 and harboring the CD4-binding site D368R mutation. This recombinant protein can interact with CoRBS and Cluster A Abs but not with CD4(57, 58)(Fig. S3). Pre-incubation of CoRBS Ab-CD4 or Cluster A Ab-CD4 with this probe significantly reduced their capacity to neutralize HIV-1 and to recognize HIV-1-infected primary CD4+ T cells (Fig. 6b,c). Conversely, competition with an inner domain stabilized gp120 probe (ID2), able to interact with Cluster A Abs (42), but not with CoRBS Abs or CD4 (Fig.S3), only affected the biological activities of N5-i5-CD4 and A32-CD4 (Fig. 6b,c). This suggests that competing with the Ab moiety significantly reduces the capacity of developed Ab-CD4 to neutralize HIV-1 and to target infected cells. All together, these results suggest that the biological activities mediated by the Ab-CD4 depend on both antibody and CD4 moiety interaction with HIV-1 Env.

**Fig. 6.**
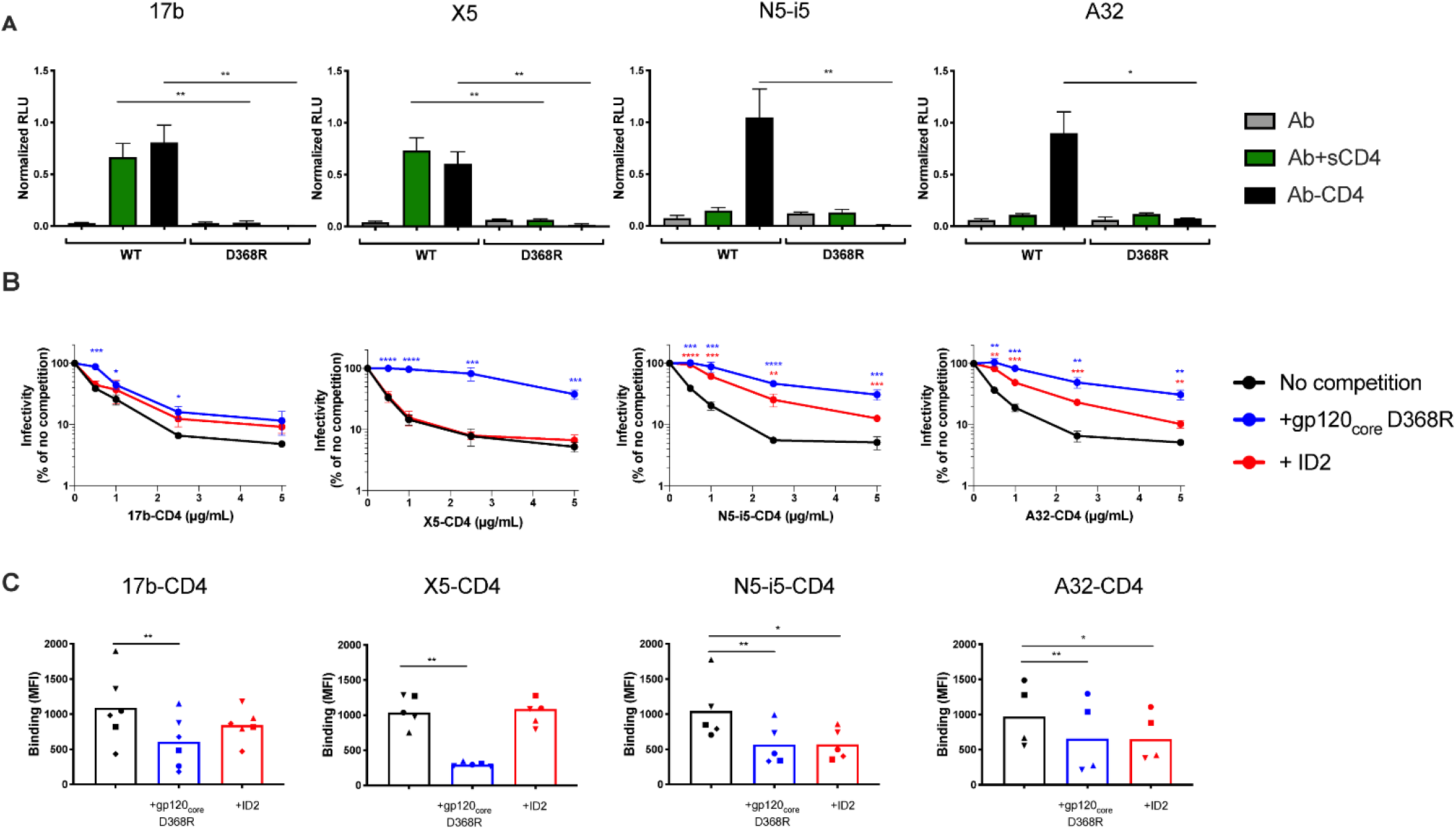
Both CD4 and antibody moieties are required to allow Ab-CD4s to target trimeric Env present on virions and infected cells. (**A**) Recognition of cellular-expressed trimeric WT or D368R Env by indicated Ab-CD4 or Ab in the presence or absence of sCD4 (3 μg/ml) was evaluated by cell-based ELISA. Data shown represent mean RLU values ± SEM from at least 3 independent experiments performed in quadruplicate, with the signal obtained from cells transfected with an empty pcDNA3.1 plasmid (no Env) subtracted, normalized to Env levels as determined by bNAb 2G12. (b-c) Ab-CD4s were pre-incubated or not with (10 μg/ml) of indicated gp120 proteins. (**B**) Viral particles pseudotyped with HIV-1 _JRFL_ Env were incubated with serial dilution of these Ab-CD4s before infecting TZM-bl cells. Luciferase activity in cell lysates was measured to determine the infectivity. Infectivity was normalized to 100% in the absence of ligand and competition. Data shown are representative of 3 independent experiments. (**C**) Recognition of primary CD4+ T cells infected with JRFL by Ab-CD4 pre-incubated or not with gp120 proteins. The graphs shown represent the compiled median fluorescence intensities on the infected (p24+) cell population. Error bars indicate means±SEM for at least 4 independent experiments. Statistical significance was tested using (A-B) unpaired or (C) paired t-test (*, *P* < 0.05; **, *P* < 0.01, ***, *P* < 0.001; ****, *P* < 0.0001)

## Discussion

Conformational CD4i gp120 epitopes within the CoRBS and C1C2 Cluster A map to conserved regions of HIV-1 Env(12, 13, 18, 19, 42, 59). Whereas the CoRBS is used as a second attachment point by the virus to specifically engage the co-receptor, the Cluster A region maps to the interior of HIV-1 trimer and is directly involved in inter-protomer contacts(15, 18, 42). The high conservation of these two sites results directly from their functional role; the CoRBS serves as the co-receptor attachment site and Cluster A is at the gp41-gp120 interface of the trimer in its ‘closed’ conformation that contributes to structural integrity(22). These critical functional importance of these sites for viral entry might explain why the virus maintains them occluded for Ab recognition. Recent data indicate that viral resistance to these targets extend into virally infected cells that express and retain most of Env trimers on their surface in the ‘closed’ conformation. This greatly limits the potential for CD4i epitopes to be targets for antibodies capable of ADCC of infected cells(16, 17, 24, 27–29). This represents a significant drawback limiting the potential of CoRBS and Cluster A Abs as agents capable of effective infected cell elimination(21, 23, 29). This is of particular importance since Cluster A Abs are known to be very efficient mediators of ADCC towards cells exposing Env in the ‘open’, membrane CD4-bound conformation (19–21, 24, 26, 29, 52).

Structural and functional analyses indicate that CoRBS and Cluster A epitopes become available for Ab recognition sequentially upon triggering with sCD4 or CD4mc of the ‘closed’ Env trimer residing on virions or infected cells. Whereas CoRBS epitopes are ‘easily’ induced and are fully formed upon triggering of Env with sCD4 or CD4mc (21, 23, 29), the Cluster A region is not so easily exposed. Additional energy is required to further trigger the structural rearrangements of the trimer that ultimately leads to the exposure of the Cluster A region. This additional energy could be provided by membrane anchored CD4 or an Ab partner with a CoRBS specificity(17, 21, 23, 29, 34). Interestingly, the bivalency of the CoRBS Ab partner is required as Fabs failed as helpers in this process.

Here we aimed to develop a single chain molecule capable of revealing the full potential of CoRBS and Cluster A as targets for ADCC of HIV-1 infected cells and improve/activate their neutralization activity. This was of particular interest for Cluster A epitopes because these map to the most conserved regions of HIV Env(12, 18, 22, 42). They are also targeted by Abs that share similar characteristics with a moderate length CDR H3 and a low degree of V_H_ affinity maturation therefore lack unusual structural features typical for Abs involved in neutralization(12, 13, 18, 42). This is a positive feature because it might limit immunogenicity of therapeutics developed based on these Abs. To preserve Fc-effector capacity of our hybrid Ab-CD4 molecules, we used the whole IgG1 Ab of human origin (CoRBS or Cluster A specificity) linked to the d1d2 domains of human CD4. The resultant multivalent molecule consists of an IgG1 with two CD4 moieties attached through a flexible peptide linker, approximately 152 Å long to allow for the potential interaction of all 4 binding moieties to cognate targets on the Env trimer(60). Mechanistic analyses indicate engagement of CD4 and Fab moieties in Env binding and functionality of Ab-CD4s confirming that the Ab target epitopes are exposed. However, at this point, it is difficult to describe the exact mechanism by which the multivalent Ab-CD4 molecules trigger multiple Env trimers on virion or at the infected cells surface. The Ab-CD4 hybrid proteins are multivalent molecules consisting of 4 binding arms (2 Ab Fabs and 2 CD4 moieties) capable of binding a total of 6 binding sites on Env trimer “open” to expose the Ab epitope (schematically shown for 17b-CD4 in Fig. S4a). Our attempts to visualize Ab-CD4 mediated crosslinking of BG505 SOSIP trimer, a soluble Env trimer preparation, with electron microscopy (EM) were inconclusive (Fig. S4b). In this studies we used BG505 SOSIP trimer and the Ab-CD4 molecules targeting the CoRBS only. This was due to limitations of BG505 SOSIP, which is stabilized by an inter gp120-gp41 disulfide bond and proline substitutions within gp41(61). The introduced mutations don’t interfere with the CoRBS exposition but inter protomer disulfide bond might restrain the structural rearrangements of the CD4-bound trimer limiting exposure of the Cluster A region, that maps to the gp41-gp120 interface. We observed heavy precipitation of BG505 SOSIP Env when an excess of 17b-CD4 to SOSIP was used indicating multivalent crosslinking of soluble trimers. Therefore, for EM experiments we mixed BG505 SOSIP with 17b-CD4 at a 3:1 molar ratio to give a large excess of trimer to minimize multivalent 17b-CD4 binding. The complex was then purified by SEC (Fig. S5). Shown in Fig. S4 are representative EM-images of SOSIP Env trimer treated with a mixture of Ab (17b) and sCD4 and the corresponding Ab-CD4 (17b-CD4) hybrid. Extensive crosslinking and aggregation of BG505 SOSIP was observed in both preparations. This is consistent with our functional data that indicates that the CoRBS is ‘easy induced’ and becomes readily available for Ab recognition with addition of sCD4. The aggregates observed for samples treated with the mixture of 17b and sCD4 result from crosslinking through 17b IgG which binds to epitopes exposed by CD4. The scale of the observed crosslinking is similar for SOSIPs treated with the mixture of 17b and CD4 and the 17b-CD4 hybrid molecules with the slightly higher abundance of oligomers with larger size in the sample treated with hybrid molecules. The limited resolution of EM images enables a direct comparison between the conformation states of the SOSIPs and the valency of binding for both preparations. The higher resolution images or structural studies with surface bound Env trimers are required to further investigate this mechanism.

Functional studies fully validated the Ab-CD4 design, hybrid proteins consisting of either CoRBS or Cluster A Ab arms showed effective virus recognition and ADCC killing of cells infected with the primary isolate JRFL. Interestingly, the level of binding and ADCC activities of Ab-CD4 molecules of Cluster A specificity were comparable to those observed for the Ab-CD4s of CoRBS specificity indicating similar epitope exposure. This is interesting given the existing constrains related to Cluster A region exposure. Moreover, this indicates that Cluster A Ab-CD4 molecules acting through multivalent binding are capable of overcoming the energy barrier required to expose the Cluster A targets residing at the infected cell surface. Studies are underway to fully understand this mechanism and to describe the overall conformation of the Env trimer in the CoRBS Ab-CD4- and the Cluster A-CD4-bound states.

Finally, our Ab-CD4 hybrid molecules showed efficient, cross-clade neutralization of Tier 1 and 2 viruses. Whereas this observation is not surprising for CoRBS Ab-CD4 molecules, it is new and significant for the Cluster A Ab-CD4 class. Utility of the CoRBS as a plausible target for Abs with neutralizing activity was already shown by several single chain molecule approaches including a single-chain variable region construct of a CoRBS Ab linked to sCD4 or eCD4-Ig variants linking CD4-IgG1 to CoRBS specific sulfopeptides(37, 39). In contrast, the Cluster A region is known to be targeted by Abs lacking any neutralizing activity including for easy to neutralize Tier 1 viruses(19). Cluster A Ab-CD4 molecules are therefore capable of stabilizing Env in more “open” conformations, potentially prematurely unleashing the energy required to fuel viral entry. This is to our knowledge the first successful attempt to develop a hybrid protein with non-neutralizing Cluster A Abs that results in a molecule capable of potent ADCC and gaining neutralizing activity against HIV-1 using these conserved and important Env targets.

## Acknowledgments

We acknowledge the Institute of Human Virology (IHV) of University of Maryland, Baltimore, MD were this work was initiated and our colleagues Dr. George K. Lewis and Dr. Krishanu Ray of IHV for their support. The authors thank the FRQS AIDS and Infectious Disease Network and Mario Legault for cohort coordination, the BSL3 and FACS facilities at CRCHUM. We thank Dennis Burton and Malcolm Martin for the infectious molecular clone JRFL and pHIV-1_AD8_, respectively. We are thankful for subject’s participation and collaboration. This work was supported by NIH R01 AI129769 to M.P. and A.F. This work was also supported by R01 AI116274 to M.P., by R01 AI150322-01 to A.F., by P01 GM56550/AI150741 to A.F., by P01 AI120756 to Georgia Tomaras and M.P and by a CIHR foundation grant #352417 to A.F. This work was also supported by a Canada Foundation for Innovation grant #41027 to A.F. A.F. is the recipient of a Canada Research Chair on Retroviral Entry. R.G. is the recipient of a MITCS postdoctoral fellowship. J.P. is the recipient of a CIHR PhD fellowship. The funders had no role in study design, data collection and analysis, decision to publish, or preparation of the manuscript. The authors have no conflicts of interest to report.

## Disclaimer

The views expressed in this presentation are those of the authors and do not reflect the official policy or position of the Uniformed Services University, US Army, the Department of Defense, or the US Government.

## Author contributions

J.R, D.N.N, W.D.T., A.F and M.P conceived the study. J.R, D.N.N, W.D.T, R.G, S.D, D.V, S.Y.G, J.P, G.G.L, H.M, S.G, A.F and M.P generated reagents, performed experiments and analyzed the data. J.R, D.N.N, W.D.T., A.F and M.P wrote the paper. All authors contributed to editing the manuscript.

## Materials and Methods

### Ethics Statement

Written informed consent was obtained from all study participants and research adhered to the ethical guidelines of CRCHUM and was reviewed and approved by the CRCHUM institutional review board (ethics committee, approval number CE16.164-CA). Research adhered to the standards indicated by the Declaration of Helsinki. All participants were adult and provided informed written consent prior to enrolment in accordance with Institutional Review Board approval.

### Cell lines and isolation of primary cells

HEK293T human embryonic kidney cells, HeLa TZMbl cells and human osteosarcoma (HOS) cells were grown as previously described (22, 47). Expi293F™ cells (Thermo Fisher Scientific) were cultured in Gibco™ Expi293™ Expression Medium supplemented with 100 IU penicillin and 100 μg/ml streptomycin solution at 37°C, 8% CO_2_, 90% humidity and shaking at 125 rotations per minute (rpm) according to the manufacturer’s protocol. Primary human peripheral blood mononuclear cells (PBMCs) and CD4^+^ T cells were isolated, activated, and cultured as previously described (40). Briefly, PBMCs were obtained by leukapheresis and CD4^+^ T lymphocytes were purified from resting PBMCs by negative selection using immunomagnetic beads per the instructions of the manufacturer (StemCell Technologies, Vancouver, BC, Canada) and were activated with phytohemagglutinin-L (10 μg/ml) for 48 h and then maintained in RPMI 1640 complete medium supplemented with recombinant interleukin-2 (rIL-2) (100 U/ml).

### Plasmids, vectors and proviral constructs

The JR-CSF IMC and the pNL4.3 Nef- Luc Env- plasmid were obtained from the AIDS Research and Reference Reagent Program, Division of AIDS, NIAID, NIH. The JRFL and pHIV-1_AD8_ IMCs were previously described (62, 63). Plasmid expressing the full-length Envs HIV-1_JRFL_, HIV-1_YU2_, HIV-1_CM244_, HIV-1_C1086_, HIV-1_BG505_ and HIV-1_ZM109_ were previously reported (22, 64–66).The CD4-IgH plasmid consists of domains 1 and 2 of CD4 fused to the heavy chain through a flexible 40 amino acids linker (6 repeats of G_4_S and 2 repeats of G_4_T motifs). Two restriction sites NheI at the 5’ end and BamHI at the 3’ end were incorporated into the CD4-polypeptide linker plasmid which was then inserted into pACP-tag(m)-2 plasmid (New England Biolabs) to obtain pACP-CD4. The IgGH gene lacking the leader sequence was amplified by PCR with the given primers and purified using the MinElute Reaction cleanup kit (Qiagen), followed by digestion with BamHI-Hf and NotI restriction enzymes (New England Biolabs). The digested fragment was purified by agarose gel electrophoresis, then ligated into BamHI and NotI sites of pACP-CD4 to afford the respective pACP-CD4-IgH vector which was transformed into NEB® 5-alpha F’Iq Competent E. coli according to the manufacturer’s protocol (New England Biolabs). Vectors containing CD4 and heavy chain genes were then sequenced and compared with the original heavy chain sequences. In preparation for larger scale protein production, plasmids were grown under ampicillin selection and purified using GeneJET plasmid Midiprep Kit (Thermo Scientific), following the protocol specified by the manufacturer. The vesicular stomatitis virus G (VSV-G)-encoding plasmid pSVCMV-IN-VSV-G was previously reported (67).

### Generation of stable CD4-IgH cell lines

Expi293 cells were seeded at 1 × 10^6^ cells/ml (viability >90%) and transfected with CD4-IgH plasmid encoding domains 1 and 2 of CD4 fused to the N-terminus of the heavy chain of selected nnAbs using EndoFectin™ Max transfection reagent (GeneCopoeia) following the manufacturer’s protocol. Transfected cells were incubated for 5 days at 37°C, in a humidified atmosphere of 8% CO2, at 125 rpm. Then, cell culture supernatants were replaced and maintained using Gibco™ Expi293™ Expression Medium supplemented with 50 μg/ml Geneticin until cell viability reached and remained > 90%.

### Expression and purification of recombinant proteins and CD4-IgG conjugates

Soluble CD4 (sCD4) was produced and purified as previously described (22). The recombinant gp120 proteins ΔV1V2V3V5 WT, ΔV1V2V3V5 D368R and ID2 were produced and purified as previously reported(42, 57). CD4-conjugated antibodies were produced in stable cell lines via transient transfection of plasmid coding light chains using EndoFectin™ Max transfection agent (GeneCopoeia). Transfected cells were incubated at 37°C, in a humidified atmosphere of 8% CO2, on an orbital shaker (125 rpm). Five days post-transfection, cell culture supernatants were collected, centrifuged and filtered through 0.22 μm PES membrane to remove cell debris. Antibody conjugates were affinity purified by protein A affinity chromatography according to the manufacturer’s instructions. The eluted proteins were concentrated, buffer-exchanged with DPBS buffer (pH 7) and purified over Superdex 200 Increase 10/300 GL column (GE Healthcare) in DPBS buffer. The purified proteins were pooled, concentrated with Amicon Ultra-15 centrifugal filters (MWCO 30,000, Millipore), analyzed by reducing SDS-PAGE with Coomassie Blue staining, and further purified by size exclusion chromatography (Superdex 200 Increase 5/150 GL column, GE Health Sciences). Finally, protein concentration was quantified by the Pierce BCA protein assay (Thermo Scientific), aliquoted and stored at −20°C.

### Negative stain electron microscopy (NS-EM)

The SOSIP CD4i Abs complexes were prepared by mixing a 3:1 molar ratio of BG505 SOSIP.664 and CD4i Abs. The resulting mixture was incubated at room temperature for 2 h and purified by SEC on a Superdex 200 Increase 10/300 GL column. Fraction containing SOSIP-CD4Abs complexes were analyzed by negative stain EM. A 5 μL of sample were applied for 1 min to carbon-coated 400 Cu mesh grid which has been glow discharged at 25 mA for 2 min, followed by negative staining with 2% Uranyl Acetate for 1 min. Data was collected using a JEOL LEM-1011 microscope operating at 100 keV, with a magnification of 120,000x.

### Viral production and infections

VSV-G-pseudotyped HIV-1 JRFL virus was produced and titrated as previously described (21). Viruses were then used to infect activated primary CD4+ T cells from healthy HIV-1-negative donors by spin infection at 800 × *g* for 1 h in 96-well plates at 25°C. For the viral neutralization assay, TZM-bl cells were infected with either single-round luciferase-expressing HIV-1 pseudovirions or fully replicative WT viruses. Briefly, 293T cells were transfected by the calcium phosphate method with the proviral vector pNL4.3Luc Env- and a plasmid expressing indicated HIV-1 Env at a ratio of 2:1 or with full IMCs constructs. Two days after transfection, the cell supernatants were harvested. Each virus preparation was frozen and stored in aliquots at –80°C until use.

### Virus Capture Assay

The assay was modified from a previously published method(34). Briefly, pseudoviral particles were produced by transfecting 2 × 10^6^ HEK293T cells with pNL4.3 Nef- Luc Env-(3.5 μg), pSVCMV-IN-VSV-G (1μg) and plasmid (2.5 μg) encoding for JRFL full length Env using the standard calcium phosphate protocol. 48 hours later, virion-containing supernatants were collected, and the cell debris was removed through centrifugation (486× g for 10 min). To immobilize antibodies on ELISA plates, white Pierce™ protein A coated 96-well plates (Thermo Fisher Scientific, Waltham, MA, USA.) were incubated with 5 μg/ml of tested antibodies diluted in 100 μL phosphate-buffered saline (PBS) overnight at 4 °C. Unbound antibodies were removed by washing the plates twice with PBS. Plates were subsequently blocked with 3% BSA in PBS for 1 hour at room temperature. After two washes with PBS, 200 μL of virion-containing supernatant was added to the wells in the presence or absence of sCD4 (10μg/ml). Viral capture by any given antibodies was visualized by adding 1 × 10^4^ HIV-1-resistant HEK293T cells in full DMEM medium per well. Forty-eight hours post-infection, cells were lysed by the addition of 30 μL of passive lysis buffer (Promega, Madison, WI, USA.) and three freeze-thaw cycles. An LB941 TriStar luminometer (Berthold Technologies) was used to measure the luciferase activity of each well after the addition of 100 μL of luciferin buffer (15 mM MgSO_4_, 15 mM KH_2_PO_4_ (pH 7.8), 1 mM ATP, and 1 mM dithiothreitol) and 50 μL of 1 mM D-luciferin potassium salt (Prolume, Randolph, VT, USA.). Luciferase signals were then normalized to those obtained with the 2G12 antibody.

### Viral neutralization assay

TZM-bl target cells (NIH AIDS reagent program) were seeded at a density of 1×10^4^ cells/well in 96-well luminometer-compatible tissue culture plates (Perkin Elmer) 24h before infection. 100μL of recombinant viruses were mixed and incubated with 100μL of Ab (+/−sCD4) or Ab-CD4 for 1h at 37°C. For the competition experiments, Ab-CD4 were pre-incubated with 10μg/ml of recombinant gp120 proteins prior incubation with recombinant virus. Each mix of virions and Ab or Ab-CD4 was then split into two and added to the target cells followed by incubation for 4h at 37°C; 100μL of fresh DMEM 5% FCS 1%Pen-Strep was then added to the cells, which have been incubated for an additional 48h at 37°C. Cells were then lysed by the addition of 30μl of passive lysis buffer (Promega) followed by one freeze-thaw cycle. An LB941 TriStar luminometer (Berthold Technologies) was used to measure the luciferase activity of each well after the addition of 100μl of luciferin buffer (15mM MgSO_4_, 15mM KPO_4_ [pH 7.8], 1mM ATP, and 1mM dithiothreitol) and 50μl of 1mM d-luciferin potassium salt (Prolume). The neutralization half-maximal inhibitory concentration (IC_50_) has been calculated with Graphpad Prism version 8.0.1.

### ELISA

The capacity of recombinant gp120 proteins to interact with CD4, CoRBS and Cluster A Abs was tested by ELISA as previously decribed(68). Bovine serum albumin (BSA) and the recombinant gp120 proteins (ΔV1V2V3V5 WT, ΔV1V2V3V5 D368R and ID2) were prepared in PBS (0.1 μg/mL) and adsorbed to MaxiSorp; Nunc plates (Thermo Fisher Scientific, Waltham, MA, USA) overnight at 4 °C. BSA was used as a negative control. Coated wells were subsequently blocked with blocking buffer (Tris-buffered saline (TBS) containing 0.1% Tween 20 and 2% [wt/vol] BSA) for 90 min at room temperature. Wells were then washed 4 times with washing buffer (Tris-buffered saline [TBS] containing 0.1% Tween 20). Abs (17b, N5i5 or CD4-Ig) were diluted in blocking buffer and incubated for 120 min at room temperature. Wells were then washed 4 times with washing buffer. This was followed by incubation of HRP conjugated antibody specific for the Fc region of human IgG (Pierce) for 90 min at room temperature. Wells were then washed 4 times with washing buffer. HRP enzyme activity was determined after the addition of a 1:1 mix of Western Lightning ECL reagents (Perkin Elmer Life Sciences, Waltham, MA, USA). Light emission was measured with an LB 941 TriStar luminometer (Berthold Technologies, Bad Wildbad, Germany).

### Cell-based ELISA

Recognition of trimeric HIV-1_JRFL_ envelope glycoprotein (Env) ΔCT at the surface of HOS cells was performed by cell-based ELISA, as previously described(48). Briefly, HOS cells were seeded in T-25 flasks (2 × 10^6^ cells per flask) and transfected the next day with a total of 12 μg of pcDNA3.1 expressing the codon-optimized HIV-1_JRFL_ EnvΔCT, either WT or containing the CD4-binding site D368R mutation, per flask using the standard polyethylenimine (PEI; Polysciences Inc., PA, USA) transfection method. Twenty-four hours after transfection, cells were plated in 384-wells plates (2×10^4^ cells per well). One day later, cells were incubated in blocking buffer (washing buffer [25 mM Tris (pH 7.5), 1.8 mM CaCl_2_, 1.0 mM MgCl_2_ and 140 mM NaCl] supplemented with 10 mg/ml non-fat dry milk and 5 mM Tris [pH 8.0] for 30 min and then co-incubated for 1 h with indicated Ab-CD4 or unconjugated Ab (1μg/ml) in the presence or absence of soluble CD4 (sCD4) (3 μg/ml) in phosphate-buffered saline [PBS] diluted in blocking buffer. Cells were then washed five times with blocking buffer and five times with washing buffer. A HRP conjugated antibody specific for the Fc region of human IgG (Pierce) was then incubated with all the samples for 45 min and then washed again as just described. All incubations were done at room temperature. To measure the HRP enzyme activity, 20 μl of a 1:1 mix of Western Lightning oxidizing and enhanced luminol reagents (Perkin Elmer Life Sciences) was added to each well. Chemiluminescence signal was acquired for 1 sec/well with the LB 941 TriStar luminometer (Berthold Technologies).

### Flow cytometry analysis of cell surface staining

Cell surface staining of infected cells was performed as previously described (40). Binding of cell surface HIV-1 Env by Ab-CD4 or unconjugated Ab (10 μg/ml) was performed at 48 h postinfection in the presence or absence of sCD4 (10 μg/ml). For competition experiments, Ab or Ab-CD4 were pre-incubated 30 min at room temperature with purified soluble gp120 ΔV1V2V3V5 D368R or ID2 protein (10 μg/ml) prior incubation with infected cells for 45min. Cells were then washed twice with PBS and stained with 2 μg/ml goat anti-human (Alexa Fluor 647; Invitrogen) secondary Abs for 15 min in PBS. After two more PBS washing, cells were fixed in a 2% PBS-formaldehyde solution. To evaluate F240 epitope exposure, infected cells were stained with Alexa-Fluor 647-conjugated F240 Abs in the presence of Ab-CD4 or unconjugated Ab (10 μg/ml) +/− sCD4 (10 μg/ml). Infected cells were then stained intracellularly for HIV-1 p24, using a Cytofix/Cytoperm fixation/permeabilization kit (BD Biosciences, Mississauga, ON, Canada) and fluorescent anti-p24 MAb (phycoerythrin [PE]-conjugated anti-p24, clone KC57; Beckman Coulter/Immunotech). The percentage of infected cells (p24^+^) was determined by gating the living cell population on the basis of viability dye staining (Aquavivid; Thermo Fisher Scientific). Samples were acquired on an LSR II cytometer (BD Biosciences), and data analysis was performed using FlowJo vX.0.7 (Tree Star, Ashland, OR, USA).

### FACS-based ADCC assay

Measurement of ADCC using the FACS-based assay was performed at 48 h post-infection as previously described(47). Briefly, infected primary CD4^+^ T cells were stained with Aquavivid viability dye and cell proliferation dye (eFluor670; eBioscience) and used as target cells. Autologous PBMC effector cells, stained with another cellular marker (cell proliferation dye eFluor450; eBioscience), were added at an effector/target ratio of 10:1 in 96-well V-bottom plates (Corning, Corning, NY). Antibodies (+/− sCD4) or Ab-CD4 were added to appropriate wells, and the cells were incubated for 15 min at room temperature. The plates were subsequently centrifuged for 1 min at 300 × *g* and incubated at 37°C and 5% CO_2_ for 5 h before being fixed in a 2% PBS–formaldehyde solution. Samples were acquired on an LSR II cytometer (BD Biosciences), and data analysis was performed using FlowJo vX.0.7 (Tree Star). The percentage of ADCC resulting from gating performed on infected lived target cells was calculated with the following formula: (percentage of p24^+^ cells in targets plus effectors) − (percentage of p24^+^ cells in targets plus effectors plus Abs)/(percentage of p24^+^ cells in targets).

### Statistical analysis

Statistics were analyzed using GraphPad Prism version 8.4.3 (GraphPad, San Diego, CA, USA). Every data set was tested for statistical normality, and this information was used to apply the appropriate (parametric or nonparametric) statistical test. *P* values of <0.05 were considered significant, and significance values are indicated as follows: *, *P* < 0.05; **, *P* < 0.01; ***, *P* < 0.001; ****, *P* < 0.0001.

---

**Fig. S1.**
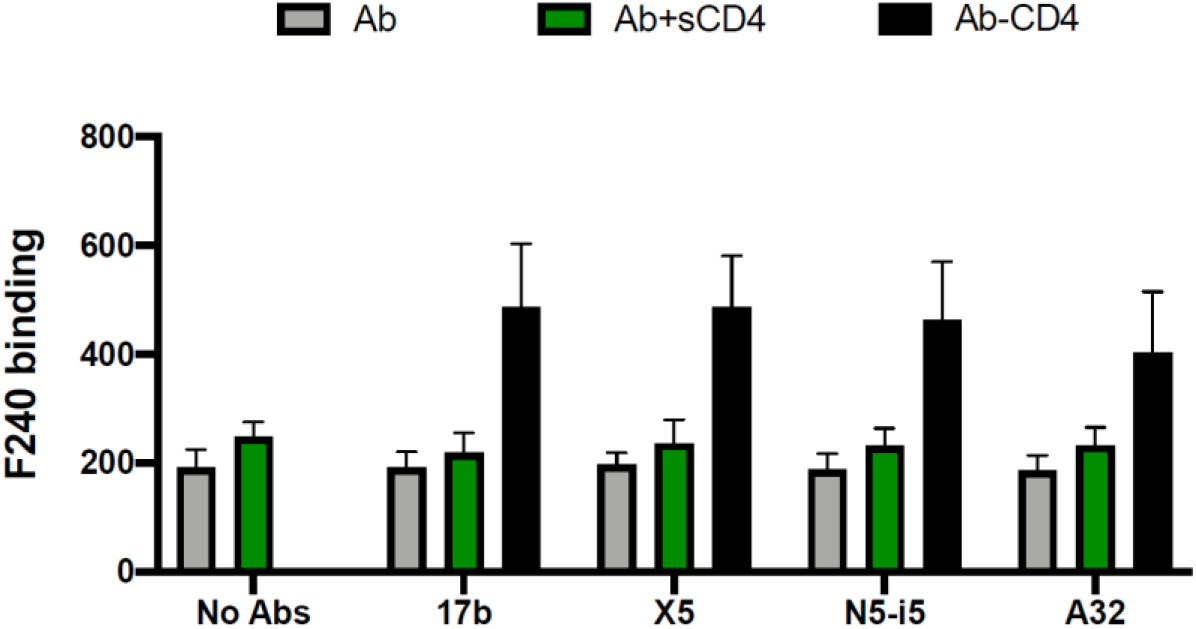
Ab-CD4s enhance F240 epitope exposure on HIV-1-infected cells. Recognition of primary CD4+ T cells infected with HIV-1 JRFL by Alexa-Fluor 647-conjugated F240 mAbs in the presence of Ab-CD4 or Ab (10 μg/ml) +/− sCD4 (10 μg/ml) was evaluated by flow cytometry. The graph shown represents the compiled median fluorescence intensities on the infected (p24+) cell population. Error bars indicate means±SEM for at 3 independent experiments.

**Fig. S2.**
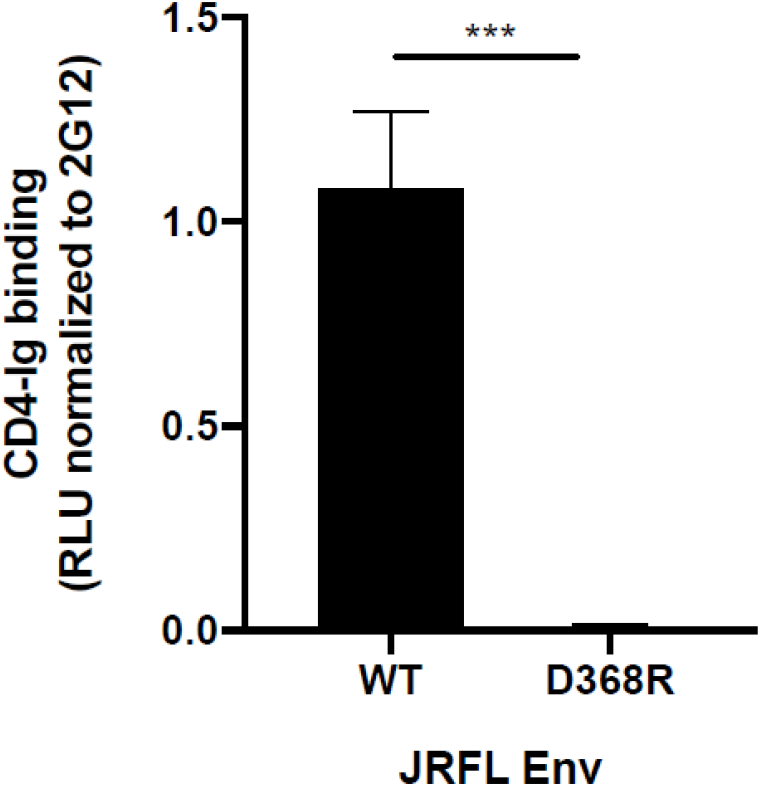
Introducing the D368R mutation in Env prevents CD4-Env interaction. The capacity of CD4-Ig to interact with either WT or D368R JRFL Env was evaluated by Cell-based ELISA. Data shown represent mean RLU values ± SEM from 5 independent experiments performed in quadruplicate, with the signal obtained from cells transfected with an empty pcDNA3.1 plasmid (no Env) subtracted, normalized to Env levels as determined by bNAb 2G12.

**Fig. S3.**
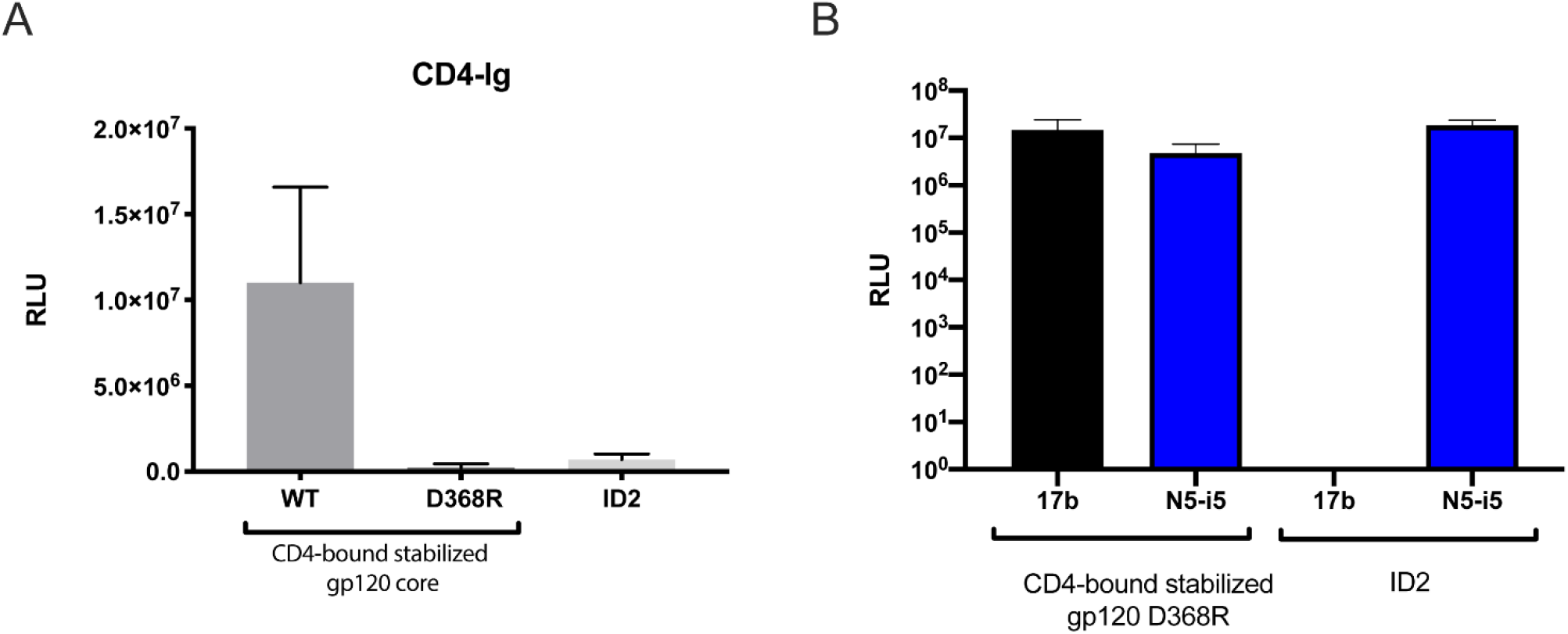
Interaction of the gp120 probes with CD4, CoRBS and Cluster A antibodies. The capacity of the recombinant gp120 proteins ΔV1V2V3V5 WT, ΔV1V2V3V5 D368R and ID2 to interact with (**A**) CD4-Ig and the (**B**) CD4-induced nnAbs (17b and N5-i5) was evaluated by standard ELISA. Data shown represent mean RLU values ± SEM from 2 independent experiments performed in triplicate, with the signal obtained with BSA subtracted.

**Fig. S4.**
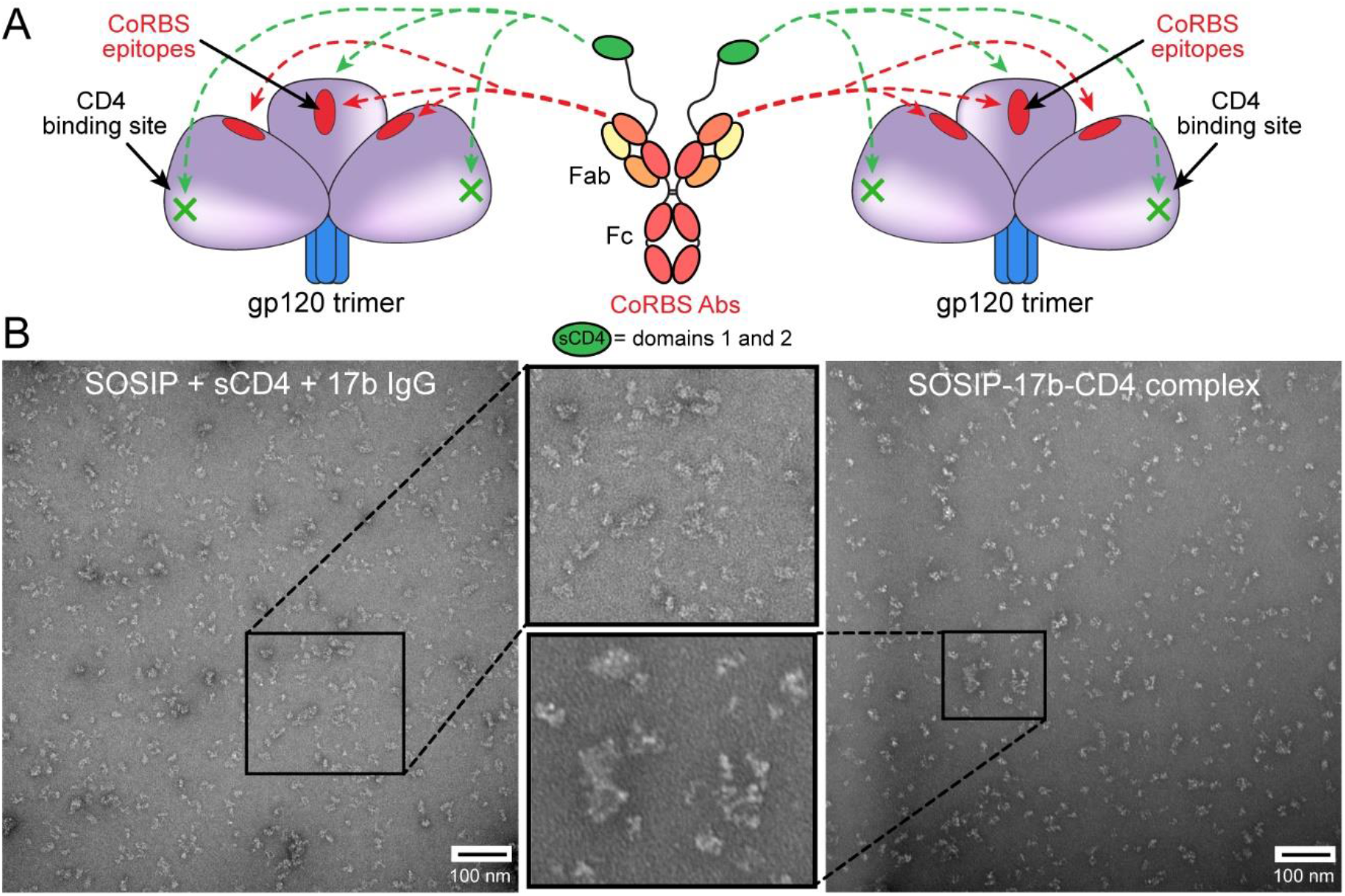
Visualization of SOSIP Ab-CD4 complexes by negative stain EM. (**A**) A cartoon illustration of the triggered and exposed CoRBS epitope depicted on an “open” trimer. Red dash arrows indicate possible binding sites of CoRBS Ab. Green dash arrows indicate possible binding site for sCD4. (**B**) Left image: raw negative stain EM of a mixture of SOSIP, CD4 and 17b IgG. Right image: raw negative stain EM of SOSIP-17b-CD4 complex. Both negative stain images are at 120,000x magnification. Middle panel are enlarged images of the selected area in order to visualize particles. The scale bar represents 100 nm.

**Fig. S5.**
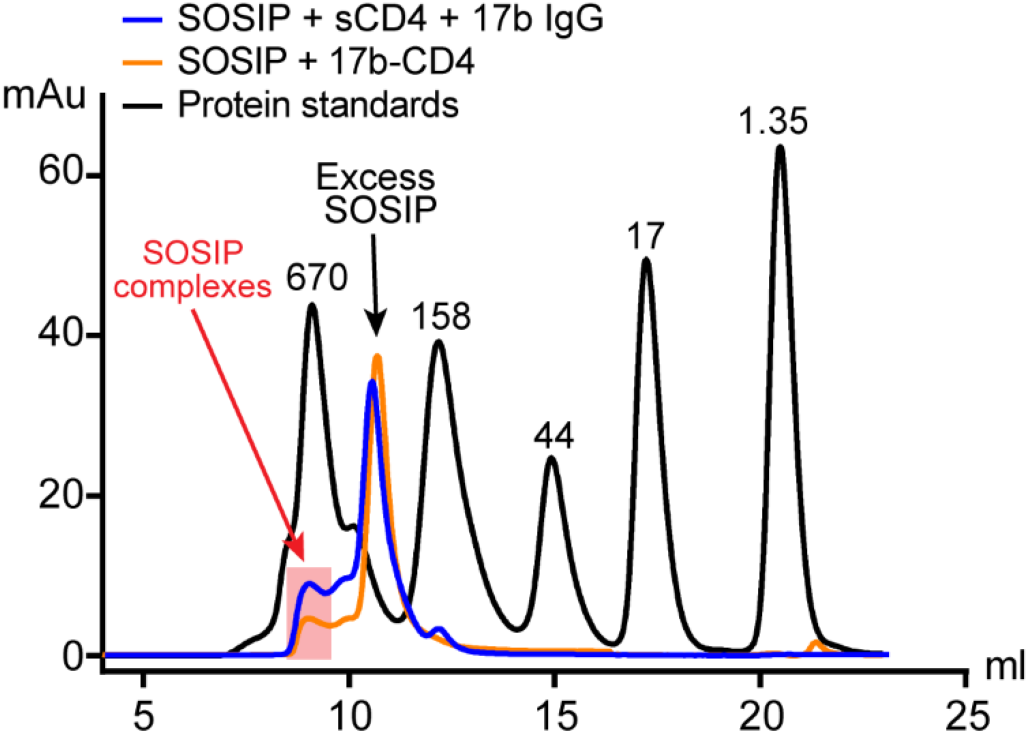
Size exclusion chromatography (SEC) profiles of SOSIP complexes. The UV chromatography of BG505 SOSIP in complex with either CD4 and 17b IgG1 (blue) or CD4-17b fusion molecule (orange) using Superdex 200 10/300 GL column. SOSIP complexes highlighted in a red box were collected and visualized by negative stain EM without concentration.

## References

1. Kowalski M, Potz J, Basiripour L, Dorfman T, Goh WC, Terwilliger E, Dayton A, Rosen C, Haseltine W, Sodroski J. 1987. Functional regions of the envelope glycoprotein of human immunodeficiency virus type 1. Science 237:1351–5.

2. Alkhatib G, Combadiere C, Broder CC, Feng Y, Kennedy PE, Murphy PM, Berger EA. 1996. CC CKR5: a RANTES, MIP-1alpha, MIP-1beta receptor as a fusion cofactor for macrophage-tropic HIV-1. Science 272:1955–8.

3. Choe H, Farzan M, Sun Y, Sullivan N, Rollins B, Ponath PD, Wu L, Mackay CR, LaRosa G, Newman W, Gerard N, Gerard C, Sodroski J. 1996. The beta-chemokine receptors CCR3 and CCR5 facilitate infection by primary HIV-1 isolates. Cell 85:1135–48.

4. Deng H, Liu R, Ellmeier W, Choe S, Unutmaz D, Burkhart M, Di Marzio P, Marmon S, Sutton RE, Hill CM, Davis CB, Peiper SC, Schall TJ, Littman DR, Landau NR. 1996. Identification of a major co-receptor for primary isolates of HIV-1. Nature 381:661–6.

5. Doranz BJ, Rucker J, Yi Y, Smyth RJ, Samson M, Peiper SC, Parmentier M, Collman RG, Doms RW. 1996. A dual-tropic primary HIV-1 isolate that uses fusin and the beta-chemokine receptors CKR-5, CKR-3, and CKR-2b as fusion cofactors. Cell 85:1149–58.

6. Dragic T, Litwin V, Allaway GP, Martin SR, Huang Y, Nagashima KA, Cayanan C, Maddon PJ, Koup RA, Moore JP, Paxton WA. 1996. HIV-1 entry into CD4+ cells is mediated by the chemokine receptor CC-CKR-5. Nature 381:667–73.

7. Feng Y, Broder CC, Kennedy PE, Berger EA. 1996. HIV-1 entry cofactor: functional cDNA cloning of a seven-transmembrane, G protein-coupled receptor. Science 272:872–7.

8. Trkola A, Dragic T, Arthos J, Binley JM, Olson WC, Allaway GP, Cheng-Mayer C, Robinson J, Maddon PJ, Moore JP. 1996. CD4-dependent, antibody-sensitive interactions between HIV-1 and its co-receptor CCR-5. Nature 384:184–7.

9. Wu L, Gerard NP, Wyatt R, Choe H, Parolin C, Ruffing N, Borsetti A, Cardoso AA, Desjardin E, Newman W, Gerard C, Sodroski J. 1996. CD4-induced interaction of primary HIV-1 gp120 glycoproteins with the chemokine receptor CCR-5. Nature 384:179–83.

10. Lu M, Blacklow SC, Kim PS. 1995. A trimeric structural domain of the HIV-1 transmembrane glycoprotein. Nat Struct Biol 2:1075–82.

11. Weissenhorn W, Dessen A, Harrison SC, Skehel JJ, Wiley DC. 1997. Atomic structure of the ectodomain from HIV-1 gp41. Nature 387:426–30.

12. Tolbert WD, Sherburn RT, Van V, Pazgier M. 2019. Structural Basis for Epitopes in the gp120 Cluster A Region that Invokes Potent Effector Cell Activity. Viruses 11.

13. Gohain N, Tolbert WD, Acharya P, Yu L, Liu T, Zhao P, Orlandi C, Visciano ML, Kamin-Lewis R, Sajadi MM, Martin L, Robinson JE, Kwong PD, DeVico AL, Ray K, Lewis GK, Pazgier M. 2015. Cocrystal Structures of Antibody N60-i3 and Antibody JR4 in Complex with gp120 Define More Cluster A Epitopes Involved in Effective Antibody-Dependent Effector Function against HIV-1. J Virol 89:8840–54.

14. Munro JB, Gorman J, Ma X, Zhou Z, Arthos J, Burton DR, Koff WC, Courter JR, Smith AB, 3rd, Kwong PD, Blanchard SC, Mothes W. 2014. Conformational dynamics of single HIV-1 envelope trimers on the surface of native virions. Science 346:759–63.

15. Tolbert WD, Gohain N, Alsahafi N, Van V, Orlandi C, Ding S, Martin L, Finzi A, Lewis GK, Ray K, Pazgier M. 2017. Targeting the Late Stage of HIV-1 Entry for Antibody-Dependent Cellular Cytotoxicity: Structural Basis for Env Epitopes in the C11 Region. Structure 25:1719–1731 e4.

16. Veillette M, Coutu M, Richard J, Batraville LA, Dagher O, Bernard N, Tremblay C, Kaufmann DE, Roger M, Finzi A. 2015. The HIV-1 gp120 CD4-Bound Conformation Is Preferentially Targeted by Antibody-Dependent Cellular Cytotoxicity-Mediating Antibodies in Sera from HIV-1-Infected Individuals. J Virol 89:545–51.

17. Veillette M, Desormeaux A, Medjahed H, Gharsallah NE, Coutu M, Baalwa J, Guan Y, Lewis G, Ferrari G, Hahn BH, Haynes BF, Robinson JE, Kaufmann DE, Bonsignori M, Sodroski J, Finzi A. 2014. Interaction with cellular CD4 exposes HIV-1 envelope epitopes targeted by antibody-dependent cell-mediated cytotoxicity. J Virol 88:2633–44.

18. Acharya P, Tolbert WD, Gohain N, Wu X, Yu L, Liu T, Huang W, Huang CC, Kwon YD, Louder RK, Luongo TS, McLellan JS, Pancera M, Yang Y, Zhang B, Flinko R, Foulke JS, Jr., Sajadi MM, Kamin-Lewis R, Robinson JE, Martin L, Kwong PD, Guan Y, DeVico AL, Lewis GK, Pazgier M. 2014. Structural definition of an antibody-dependent cellular cytotoxicity response implicated in reduced risk for HIV-1 infection. J Virol 88:12895–906.

19. Guan Y, Pazgier M, Sajadi MM, Kamin-Lewis R, Al-Darmarki S, Flinko R, Lovo E, Wu X, Robinson JE, Seaman MS, Fouts TR, Gallo RC, DeVico AL, Lewis GK. 2013. Diverse specificity and effector function among human antibodies to HIV-1 envelope glycoprotein epitopes exposed by CD4 binding. Proc Natl Acad Sci U S A 110:E69–78.

20. Ferrari G, Pollara J, Kozink D, Harms T, Drinker M, Freel S, Moody MA, Alam SM, Tomaras GD, Ochsenbauer C, Kappes JC, Shaw GM, Hoxie JA, Robinson JE, Haynes BF. 2011. An HIV-1 gp120 envelope human monoclonal antibody that recognizes a C1 conformational epitope mediates potent antibody-dependent cellular cytotoxicity (ADCC) activity and defines a common ADCC epitope in human HIV-1 serum. J Virol 85:7029–36.

21. Alsahafi N, Bakouche N, Kazemi M, Richard J, Ding S, Bhattacharyya S, Das D, Anand SP, Prevost J, Tolbert WD, Lu H, Medjahed H, Gendron-Lepage G, Ortega Delgado GG, Kirk S, Melillo B, Mothes W, Sodroski J, Smith AB, 3rd, Kaufmann DE, Wu X, Pazgier M, Rouiller I, Finzi A, Munro JB. 2019. An Asymmetric Opening of HIV-1 Envelope Mediates Antibody-Dependent Cellular Cytotoxicity. Cell Host Microbe 25:578–587 e5.

22. Finzi A, Xiang SH, Pacheco B, Wang L, Haight J, Kassa A, Danek B, Pancera M, Kwong PD, Sodroski J. 2010. Topological layers in the HIV-1 gp120 inner domain regulate gp41 interaction and CD4-triggered conformational transitions. Mol Cell 37:656–67.

23. Richard J, Pacheco B, Gohain N, Veillette M, Ding S, Alsahafi N, Tolbert WD, Prevost J, Chapleau JP, Coutu M, Jia M, Brassard N, Park J, Courter JR, Melillo B, Martin L, Tremblay C, Hahn BH, Kaufmann DE, Wu X, Smith AB, 3rd, Sodroski J, Pazgier M, Finzi A. 2016. Co-receptor Binding Site Antibodies Enable CD4-Mimetics to Expose Conserved Anti-cluster A ADCC Epitopes on HIV-1 Envelope Glycoproteins. EBioMedicine 12:208–218.

24. Ding S, Veillette M, Coutu M, Prevost J, Scharf L, Bjorkman PJ, Ferrari G, Robinson JE, Sturzel C, Hahn BH, Sauter D, Kirchhoff F, Lewis GK, Pazgier M, Finzi A. 2016. A Highly Conserved Residue of the HIV-1 gp120 Inner Domain Is Important for Antibody-Dependent Cellular Cytotoxicity Responses Mediated by Anti-cluster A Antibodies. J Virol 90:2127–34.

25. Decker JM, Bibollet-Ruche F, Wei X, Wang S, Levy DN, Wang W, Delaporte E, Peeters M, Derdeyn CA, Allen S, Hunter E, Saag MS, Hoxie JA, Hahn BH, Kwong PD, Robinson JE, Shaw GM. 2005. Antigenic conservation and immunogenicity of the HIV coreceptor binding site. J Exp Med 201:1407–19.

26. Prevost J, Richard J, Ding S, Pacheco B, Charlebois R, Hahn BH, Kaufmann DE, Finzi A. 2018. Envelope glycoproteins sampling states 2/3 are susceptible to ADCC by sera from HIV-1-infected individuals. Virology 515:38–45.

27. Richard J, Prevost J, Baxter AE, von Bredow B, Ding S, Medjahed H, Delgado GG, Brassard N, Sturzel CM, Kirchhoff F, Hahn BH, Parsons MS, Kaufmann DE, Evans DT, Finzi A. 2018. Uninfected Bystander Cells Impact the Measurement of HIV-Specific Antibody-Dependent Cellular Cytotoxicity Responses. mBio 9.

28. Alsahafi N, Ding S, Richard J, Markle T, Brassard N, Walker B, Lewis GK, Kaufmann DE, Brockman MA, Finzi A. 2015. Nef Proteins from HIV-1 Elite Controllers Are Inefficient at Preventing Antibody-Dependent Cellular Cytotoxicity. J Virol 90:2993–3002.

29. Anand SP, Prevost J, Baril S, Richard J, Medjahed H, Chapleau JP, Tolbert WD, Kirk S, Smith AB, 3rd, Wines BD, Kent SJ, Hogarth PM, Parsons MS, Pazgier M, Finzi A. 2019. Two Families of Env Antibodies Efficiently Engage Fc-Gamma Receptors and Eliminate HIV-1-Infected Cells. J Virol 93.

30. von Bredow B, Arias JF, Heyer LN, Gardner MR, Farzan M, Rakasz EG, Evans DT. 2015. Envelope Glycoprotein Internalization Protects Human and Simian Immunodeficiency Virus-Infected Cells from Antibody-Dependent Cell-Mediated Cytotoxicity. J Virol 89:10648–55.

31. Anand SP, Grover JR, Tolbert WD, Prevost J, Richard J, Ding S, Baril S, Medjahed H, Evans DT, Pazgier M, Mothes W, Finzi A. 2019. Antibody-Induced Internalization of HIV-1 Env Proteins Limits Surface Expression of the Closed Conformation of Env. J Virol 93.

32. Arias JF, Heyer LN, von Bredow B, Weisgrau KL, Moldt B, Burton DR, Rakasz EG, Evans DT. 2014. Tetherin antagonism by Vpu protects HIV-infected cells from antibody-dependent cell-mediated cytotoxicity. Proc Natl Acad Sci U S A 111:6425–30.

33. Alvarez RA, Hamlin RE, Monroe A, Moldt B, Hotta MT, Rodriguez Caprio G, Fierer DS, Simon V, Chen BK. 2014. HIV-1 Vpu antagonism of tetherin inhibits antibody-dependent cellular cytotoxic responses by natural killer cells. J Virol 88:6031–46.

34. Ding S, Gasser R, Gendron-Lepage G, Medjahed H, Tolbert WD, Sodroski J, Pazgier M, Finzi A. 2019. CD4 Incorporation into HIV-1 Viral Particles Exposes Envelope Epitopes Recognized by CD4-Induced Antibodies. J Virol 93.

35. Laumaea A, Smith AB, 3rd, Sodroski J, Finzi A. 2020. Opening the HIV envelope: potential of CD4 mimics as multifunctional HIV entry inhibitors. Curr Opin HIV AIDS 15:300–308.

36. Madani N, Princiotto AM, Schon A, LaLonde J, Feng Y, Freire E, Park J, Courter JR, Jones DM, Robinson J, Liao HX, Moody MA, Permar S, Haynes B, Smith AB, 3rd, Wyatt R, Sodroski J. 2014. CD4-mimetic small molecules sensitize human immunodeficiency virus to vaccine-elicited antibodies. J Virol 88:6542–55.

37. Lagenaur LA, Villarroel VA, Bundoc V, Dey B, Berger EA. 2010. sCD4-17b bifunctional protein: extremely broad and potent neutralization of HIV-1 Env pseudotyped viruses from genetically diverse primary isolates. Retrovirology 7:11.

38. Dey B, Del Castillo CS, Berger EA. 2003. Neutralization of human immunodeficiency virus type 1 by sCD4-17b, a single-chain chimeric protein, based on sequential interaction of gp120 with CD4 and coreceptor. J Virol 77:2859–65.

39. Gardner MR, Kattenhorn LM, Kondur HR, von Schaewen M, Dorfman T, Chiang JJ, Haworth KG, Decker JM, Alpert MD, Bailey CC, Neale ES, Jr., Fellinger CH, Joshi VR, Fuchs SP, Martinez-Navio JM, Quinlan BD, Yao AY, Mouquet H, Gorman J, Zhang B, Poignard P, Nussenzweig MC, Burton DR, Kwong PD, Piatak M, Jr., Lifson JD, Gao G, Desrosiers RC, Evans DT, Hahn BH, Ploss A, Cannon PM, Seaman MS, Farzan M. 2015. AAV-expressed eCD4-Ig provides durable protection from multiple SHIV challenges. Nature 519:87–91.

40. Richard J, Veillette M, Brassard N, Iyer SS, Roger M, Martin L, Pazgier M, Schon A, Freire E, Routy JP, Smith AB, 3rd, Park J, Jones DM, Courter JR, Melillo BN, Kaufmann DE, Hahn BH, Permar SR, Haynes BF, Madani N, Sodroski JG, Finzi A. 2015. CD4 mimetics sensitize HIV-1-infected cells to ADCC. Proc Natl Acad Sci U S A 112:E2687–94.

41. Vezina D, Gong SY, Tolbert WD, Ding S, Nguyen D, Richard J, Gendron-Lepage G, Melillo B, 3rd, Smith AB, Pazgier M, Finzi A. 2020. Stabilizing the HIV-1 envelope glycoprotein State 2A conformation. J Virol doi:10.1128/JVI.01620-20.

42. Tolbert WD, Gohain N, Veillette M, Chapleau JP, Orlandi C, Visciano ML, Ebadi M, DeVico AL, Fouts TR, Finzi A, Lewis GK, Pazgier M. 2016. Paring Down HIV Env: Design and Crystal Structure of a Stabilized Inner Domain of HIV-1 gp120 Displaying a Major ADCC Target of the A32 Region. Structure 24:697–709.

43. Thali M, Moore JP, Furman C, Charles M, Ho DD, Robinson J, Sodroski J. 1993. Characterization of conserved human immunodeficiency virus type 1 gp120 neutralization epitopes exposed upon gp120-CD4 binding. J Virol 67:3978–88.

44. Kwong PD, Wyatt R, Robinson J, Sweet RW, Sodroski J, Hendrickson WA. 1998. Structure of an HIV gp120 envelope glycoprotein in complex with the CD4 receptor and a neutralizing human antibody. Nature 393:648–59.

45. Darbha R, Phogat S, Labrijn AF, Shu Y, Gu Y, Andrykovitch M, Zhang MY, Pantophlet R, Martin L, Vita C, Burton DR, Dimitrov DS, Ji X. 2004. Crystal structure of the broadly cross-reactive HIV-1-neutralizing Fab X5 and fine mapping of its epitope. Biochemistry 43:1410–7.

46. Moulard M, Phogat SK, Shu Y, Labrijn AF, Xiao X, Binley JM, Zhang MY, Sidorov IA, Broder CC, Robinson J, Parren PW, Burton DR, Dimitrov DS. 2002. Broadly cross-reactive HIV-1-neutralizing human monoclonal Fab selected for binding to gp120-CD4-CCR5 complexes. Proc Natl Acad Sci U S A 99:6913–8.

47. Veillette M, Coutu M, Richard J, Batraville LA, Desormeaux A, Roger M, Finzi A. 2014. Conformational evaluation of HIV-1 trimeric envelope glycoproteins using a cell-based ELISA assay. J Vis Exp doi:10.3791/51995:51995.

48. Ding S, Grenier MC, Tolbert WD, Vezina D, Sherburn R, Richard J, Prevost J, Chapleau JP, Gendron-Lepage G, Medjahed H, Abrams C, Sodroski J, Pazgier M, Smith AB, 3rd, Finzi A. 2019. A New Family of Small-Molecule CD4-Mimetic Compounds Contacts Highly Conserved Aspartic Acid 368 of HIV-1 gp120 and Mediates Antibody-Dependent Cellular Cytotoxicity. J Virol 93.

49. Mengistu M, Ray K, Lewis GK, DeVico AL. 2015. Antigenic properties of the human immunodeficiency virus envelope glycoprotein gp120 on virions bound to target cells. PLoS Pathog 11:e1004772.

50. Richard J, Veillette M, Ding S, Zoubchenok D, Alsahafi N, Coutu M, Brassard N, Park J, Courter JR, Melillo B, Smith AB, 3rd, Shaw GM, Hahn BH, Sodroski J, Kaufmann DE, Finzi A. 2016. Small CD4 Mimetics Prevent HIV-1 Uninfected Bystander CD4 + T Cell Killing Mediated by Antibody-dependent Cell-mediated Cytotoxicity. EBioMedicine 3:122–34.

51. Gohain N, Tolbert WD, Orlandi C, Richard J, Ding S, Chen X, Bonsor DA, Sundberg EJ, Lu W, Ray K, Finzi A, Lewis GK, Pazgier M. 2016. Molecular basis for epitope recognition by non-neutralizing anti-gp41 antibody F240. Sci Rep 6:36685.

52. Richard J, Prevost J, Alsahafi N, Ding S, Finzi A. 2018. Impact of HIV-1 Envelope Conformation on ADCC Responses. Trends Microbiol 26:253–265.

53. Scheid JF, Mouquet H, Feldhahn N, Seaman MS, Velinzon K, Pietzsch J, Ott RG, Anthony RM, Zebroski H, Hurley A, Phogat A, Chakrabarti B, Li Y, Connors M, Pereyra F, Walker BD, Wardemann H, Ho D, Wyatt RT, Mascola JR, Ravetch JV, Nussenzweig MC. 2009. Broad diversity of neutralizing antibodies isolated from memory B cells in HIV-infected individuals. Nature 458:636–40.

54. Xiang SH, Wang L, Abreu M, Huang CC, Kwong PD, Rosenberg E, Robinson JE, Sodroski J. 2003. Epitope mapping and characterization of a novel CD4-induced human monoclonal antibody capable of neutralizing primary HIV-1 strains. Virology 315:124–34.

55. Xiang SH, Doka N, Choudhary RK, Sodroski J, Robinson JE. 2002. Characterization of CD4-induced epitopes on the HIV type 1 gp120 envelope glycoprotein recognized by neutralizing human monoclonal antibodies. AIDS Res Hum Retroviruses 18:1207–17.

56. Richard J, Veillette M, Batraville LA, Coutu M, Chapleau JP, Bonsignori M, Bernard N, Tremblay C, Roger M, Kaufmann DE, Finzi A. 2014. Flow cytometry-based assay to study HIV-1 gp120 specific antibody-dependent cellular cytotoxicity responses. J Virol Methods 208:107–14.

57. Coutu M, Finzi A. 2015. HIV-1 gp120 dimers decrease the overall affinity of gp120 preparations for CD4-induced ligands. J Virol Methods 215-216:37–44.

58. Levast B, Barblu L, Coutu M, Prevost J, Brassard N, Peres A, Stegen C, Madrenas J, Kaufmann DE, Finzi A. 2017. HIV-1 gp120 envelope glycoprotein determinants for cytokine burst in human monocytes. PLoS One 12:e0174550.

59. Tolbert WD, Sherburn R, Gohain N, Ding S, Flinko R, Orlandi C, Ray K, Finzi A, Lewis GK, Pazgier M. 2020. Defining rules governing recognition and Fc-mediated effector functions to the HIV-1 co-receptor binding site. BMC Biol 18:91.

60. Huston JS, Levinson D, Mudgett-Hunter M, Tai MS, Novotny J, Margolies MN, Ridge RJ, Bruccoleri RE, Haber E, Crea R, et al. 1988. Protein engineering of antibody binding sites: recovery of specific activity in an anti-digoxin single-chain Fv analogue produced in Escherichia coli. Proc Natl Acad Sci U S A 85:5879–83.

61. Sanders RW, Derking R, Cupo A, Julien JP, Yasmeen A, de Val N, Kim HJ, Blattner C, de la Pena AT, Korzun J, Golabek M, de Los Reyes K, Ketas TJ, van Gils MJ, King CR, Wilson IA, Ward AB, Klasse PJ, Moore JP. 2013. A next-generation cleaved, soluble HIV-1 Env trimer, BG505 SOSIP.664 gp140, expresses multiple epitopes for broadly neutralizing but not non-neutralizing antibodies. PLoS Pathog 9:e1003618.

62. O’Brien WA, Koyanagi Y, Namazie A, Zhao JQ, Diagne A, Idler K, Zack JA, Chen IS. 1990. HIV-1 tropism for mononuclear phagocytes can be determined by regions of gp120 outside the CD4-binding domain. Nature 348:69–73.

63. Theodore TS, Englund G, Buckler-White A, Buckler CE, Martin MA, Peden KW. 1996. Construction and characterization of a stable full-length macrophage-tropic HIV type 1 molecular clone that directs the production of high titers of progeny virions. AIDS Res Hum Retroviruses 12:191–4.

64. Medjahed H, Pacheco B, Desormeaux A, Sodroski J, Finzi A. 2013. The HIV-1 gp120 major variable regions modulate cold inactivation. J Virol 87:4103–11.

65. Montefiori DC, Karnasuta C, Huang Y, Ahmed H, Gilbert P, de Souza MS, McLinden R, Tovanabutra S, Laurence-Chenine A, Sanders-Buell E, Moody MA, Bonsignori M, Ochsenbauer C, Kappes J, Tang H, Greene K, Gao H, LaBranche CC, Andrews C, Polonis VR, Rerks-Ngarm S, Pitisuttithum P, Nitayaphan S, Kaewkungwal J, Self SG, Berman PW, Francis D, Sinangil F, Lee C, Tartaglia J, Robb ML, Haynes BF, Michael NL, Kim JH. 2012. Magnitude and breadth of the neutralizing antibody response in the RV144 and Vax003 HIV-1 vaccine efficacy trials. J Infect Dis 206:431–41.

66. Prevost J, Tolbert WD, Medjahed H, Sherburn RT, Madani N, Zoubchenok D, Gendron-Lepage G, Gaffney AE, Grenier MC, Kirk S, Vergara N, Han C, Mann BT, Chenine AL, Ahmed A, Chaiken I, Kirchhoff F, Hahn BH, Haim H, Abrams CF, Smith AB, 3rd, Sodroski J, Pazgier M, Finzi A. 2020. The HIV-1 Env gp120 Inner Domain Shapes the Phe43 Cavity and the CD4 Binding Site. mBio 11.

67. Lodge R, Lalonde JP, Lemay G, Cohen EA. 1997. The membrane-proximal intracytoplasmic tyrosine residue of HIV-1 envelope glycoprotein is critical for basolateral targeting of viral budding in MDCK cells. EMBO J 16:695–705.

68. Beaudoin-Bussieres G, Prevost J, Gendron-Lepage G, Melillo B, Chen J, Smith Iii AB, Pazgier M, Finzi A. 2020. Elicitation of Cluster A and Co-Receptor Binding Site Antibodies are Required to Eliminate HIV-1 Infected Cells. Microorganisms 8.

